# Resting-state Functional Connections with the Hippocampus and with the Caudate Nucleus Predict Working Memory Performance in Multiple Sclerosis

**DOI:** 10.1101/2025.10.20.683478

**Authors:** Christopher J. Cagna, Ekaterina Dobryakova, Kai Sucich, Tien T. Tong, Joshua Sandry

**Author notes:** Corresponding author: Christopher J. Cagna. Present address: Perelman School of Medicine, The University of Pennsylvania, 3400 Civic Center Boulevard, Philadelphia, PA 19104, United States.

## Abstract

Episodic memory (EM) dysfunction is a common symptom of multiple sclerosis (MS). A related process, working memory (WM), supports long-term EM and is often also impaired. Functional connectivity (FC) between the hippocampus and caudate nucleus supports EM performance. However, this connectivity’s specific influences on WM performance in MS is relatively unknown. Resting-state FC (RSFC) possesses clinical utility, including predicting cognitive impairments before they are captured by assessment. The present study tested whether RSFC with the hippocampus and caudate could predict WM performance in people with MS and in healthy individuals (HC). In a secondary analysis of 78 participants (42 MS, 36 HC), RSFC between the hippocampus, caudate nucleus, and other regions was quantified. These connections’ predictive influence on WM performance was then examined using a global WM performance measure for a subset with available neuropsychological data (26 MS, 15 HC). Across the entire sample (N=78), MS participants displayed stronger coupling between the right hippocampus and left dorsomedial prefrontal cortex, compared to HC participants. Within MS participants, stronger coupling between the left hippocampus (LHipp) and left ventral anterior cingulate (LvACC), and between the left caudate (LCaud) and right insula, were observed. Stronger decoupling between the LHipp and left supramarginal gyrus (LSMG) also emerged. Stronger LHipp-LvACC connectivity predicted worse WM performance, whereas stronger LHipp-LSMG connectivity and LCaud-RInsula connectivity each predicted better performance. The hippocampal connections were also inversely correlated. Findings identify a caudate circuit and potential hippocampal network whose aberrant, intrinsic activity could serve as neural markers for WM dysfunction in MS.

## 1. Introduction

Multiple sclerosis (MS) is an autoimmune, demyelinating, neuroinflammatory disease that impacts 2.9 million people worldwide (MS International Foundation, 2025). MS targets both white and gray matter in the central nervous system (Pitteri et al., 2021; Tsouki & Williams, 2021). White matter degradations along affected axons result in disrupted neural transmission, which in turn produces significant motor and cognitive impairments (DeLuca et al., 2020). Cognitive impairments, in particular, have been linked to significant negative impacts on everyday functioning and quality of life in people living with MS (Chiaravalloti & DeLuca, 2008). One prevalent cognitive symptom is dysfunction in episodic memory, or the memory system that allows for the recall of distinct experiences, or “episodes,” in the past (Brissart et al., 2012; Tulving, 2002). A related memory process, working memory (WM), is associated with episodic long-term memory impairment in MS (Sandry & Sumowski, 2014; Sandry, 2015). WM is the short-term storage and active manipulation of encoded stimuli (Cowan, 1995). As such, it plays an important role in the initial encoding of stimuli that are later consolidated and formed into long-term episodic memories and is an important process to consider when understanding neurocognitive change in MS.

Within the brain, the hippocampus is a key region linked to memory ability. The hippocampus displays structural and neurochemical abnormalities in MS (Geurts et al., 2007; Papadopoulos et al., 2009; Roosendaal et al., 2008), and these abnormalities are often associated with memory performance (Muhlert et al., 2014; Sumowski et al., 2016, 2017). Additionally, altered hippocampal functioning has been linked to impairments in long-term memory functions in MS, including general memory (Hulst et al., 2015; Sacco et al., 2015) and episodic memory (González Torre et al., 2017). While historically associated with declarative long-term memory, there is evidence that the hippocampus is also involved in WM processing specifically (Borders, Ranganath, Yonelinas, 2022; Kumar et al., 2016; Nee & Jonides, 2008, 2011; Leszczynski, 2011, 2015; Yonelinas et al., 2024). One study also reports its contributions to WM performance via altered theta oscillatory processes in MS (Costers et al., 2020). Interestingly, functional connections between the hippocampus and other regions traditionally associated with nondeclarative forms of memory are also important for understanding memory impairment in MS (Hulst et al., 2015). The caudate nucleus in the dorsomedial section of the striatum (a region of the basal ganglia), was historically associated with motor control, but has more recently been recognized for its cognitive functions, including procedural learning and goal-directed behavior (Grahn et al., 2008). The caudate also interacts with the hippocampus to support learning and episodic memory performance (Freedberg et al., 2022; Freedberg, 2023; Müller et al., 2018), thus making it an additional region of focus for better understanding neural mechanisms associated with WM in MS. Despite this fact, there is little research examining the roles of both of these regions, as well as their functional connections, on WM performance in MS.

Evidence for neural networks that contribute to WM is typically garnered through functional magnetic resonance imaging (fMRI) WM tasks that elicit activation and functional connectivity between these regions. While this task-based connectivity is certainly useful for this purpose, measuring functional connectivity of the brain at “rest” (i.e., one’s spontaneous, intrinsic level of connectivity, when an individual is not actively engaged in a task) may provide additional insight. Resting-state functional connectivity (RSFC) is the degree to which blood-oxygen-level-dependent (BOLD) responses in two or more regions are temporally correlated when an individual is at rest. The greater the temporal association between these BOLD signals, the greater the likelihood these regions influence each other – either directly or indirectly through the influence of a mutual region.

RSFC has been shown to serve as a useful biomarker for clinical patterns. One’s “baseline” functional connectivity serves several useful functions. For example, RSFC can identify at-risk individuals within a clinical group (Fox & Greicius, 2010; Zhang et al., 2021) and can also predict patterns of symptom progression for individuals who have developed a disease or condition (Wisch et al., 2020). RSFC is also a useful technique for identifying baseline aberrant connectivity patterns that may signal future development of cognitive symptoms or impairments, allowing for targeted, preventative intervention before symptoms develop (Fox & Greicius, 2010; Zhang et al., 2021). The clinical utility of RSFC has been reported in various clinical populations – including Alzheimer’s Disease (Lin et al., 2018), schizophrenia (Venkataraman et al., 2012), traumatic brain injury (Venkatesan et al., 2015), and MS (Azzimonti et al., 2023; Rocca et al., 2022) – further underscoring its widespread utility in linking baseline brain connectivity with future cognitive dysfunction.

Given this clinical utility of RSFC, an individual’s spontaneous hippocampal and caudal functional connectivity is a candidate metric for better understanding WM impairment in MS. In support of this, previous work has demonstrated RSFC is a valuable metric for understanding general memory impairment in MS (Hulst et al., 2015). The present study tested whether RSFC between the hippocampus, caudate, and other brain areas could predict WM performance in people with MS and in neurologically healthy individuals (HC). In a secondary analysis of an aggregate dataset comprised of three independent datasets, we identified functional connections between the hippocampus, caudate, and regions throughout the rest of the brain at rest. We then tested whether these connections, in turn, significantly predicted WM performance using a measure that we derived from available WM neuropsychological measures. Given the roles of the hippocampus and caudate in WM, we hypothesized that functional connections with these regions <u>at rest</u> should predict WM performance scores. Observing such an effect could identify potential neural signatures of WM in people with MS, allowing for earlier targeted intervention during the time window between the detection of aberrant connectivity patterns and overt displays of WM impairment during neuropsychological assessment.

## 2. Materials and methods

### 2.1. Participants

Resting-state BOLD data from a total of 83 (45 MS, 38 HC) participants from three independent datasets were examined in the current study (Cagna et al., 2022; Zuppichini & Sandry, 2018; one unpublished). Participant recruitment, behavioral data collection, and fMRI data acquisition for all three parent studies took place at Kessler Foundation (East Hanover, New Jersey, USA; West Orange, New Jersey, USA). Each study also included the same exclusion criteria. Participants were screened prior to each study’s enrollment and were excluded if they met any of the following criteria: left-handedness, diagnosis of a neurological disease other than MS (e.g., epilepsy, stroke, traumatic brain injury); significant history of alcohol abuse, substance abuse, or psychiatric illness; current diagnosis of major depressive disorder, bipolar disorder, or schizophrenia; use of steroids, benzodiazepines, or neuroleptics within the past four weeks; and presence of ferrous metal or implanted electrical devices in the body. Additionally, MS participants were excluded if they experienced an exacerbation of symptoms (i.e., a relapse, or “flare-up”) within the past four weeks. A total of 5 (3 MS, 2 HC) participants were excluded for the following reasons: excessive head motion (1 MS, 1 HC), severe image distortion (1 HC), and missing resting-state volumes (2 MS). The final sample thus comprised 78 participants – 42 MS and 36 HC. For analyses examining the influence of hippocampus and caudate RSFC on WM performance, a subset of participants with available neuropsychological data (26 MS, 15 HC) was examined. The smaller sample for these analyses was due to each of the three parent studies not containing all constituent neuropsychological measures that were used to generate our WM performance measure (detailed in the next section). We indicate analyses with this smaller subset in all figures and in degrees of freedom values for respective analyses. Demographic information for the samples appears in Table 1. MS diagnosis was verified through medical records. Each study was approved by the Institutional Review Boards of Kessler Foundation and Montclair State University. All participants provided informed consent, and all received compensation for their participation.

**Table 1.**
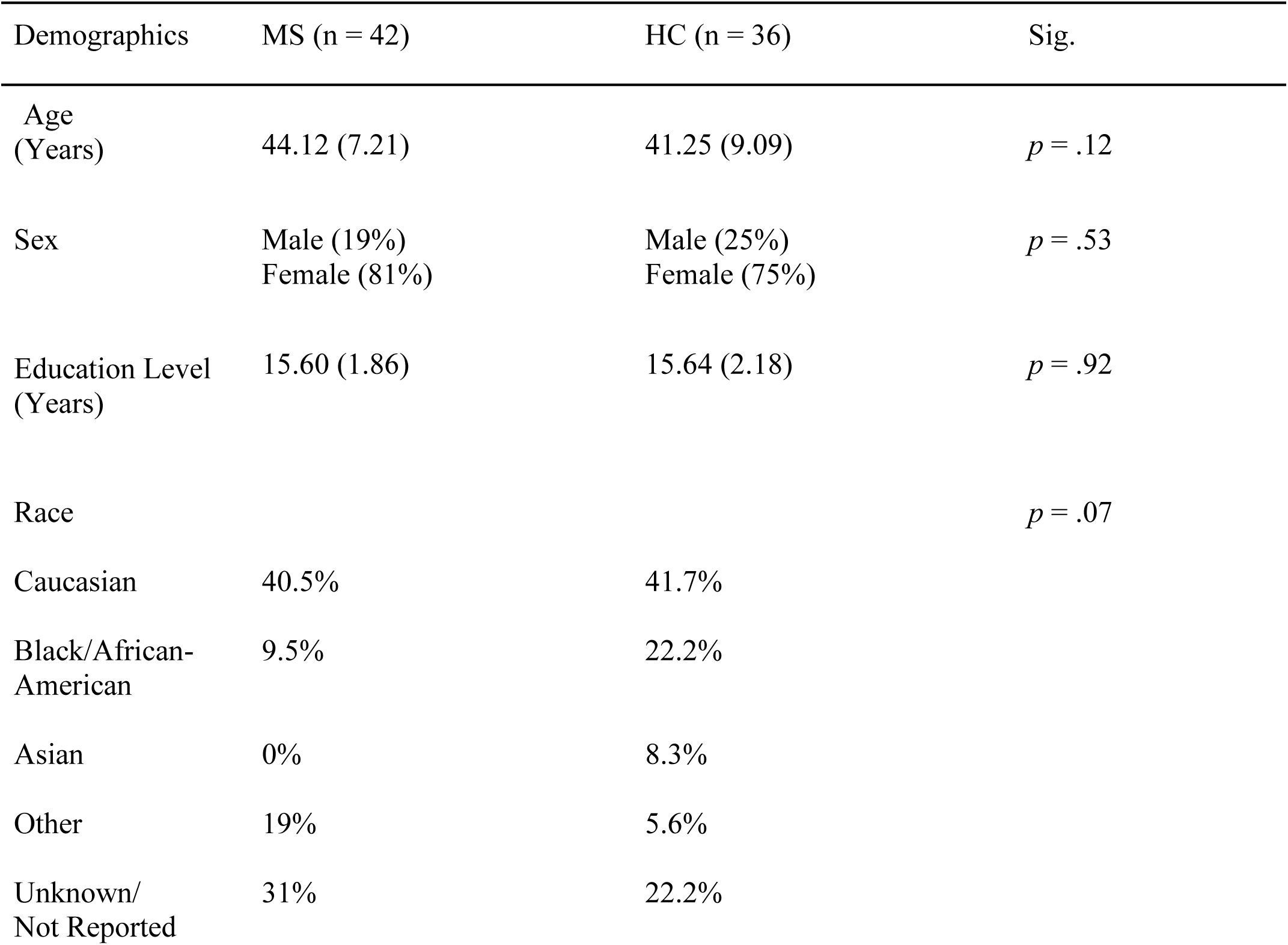
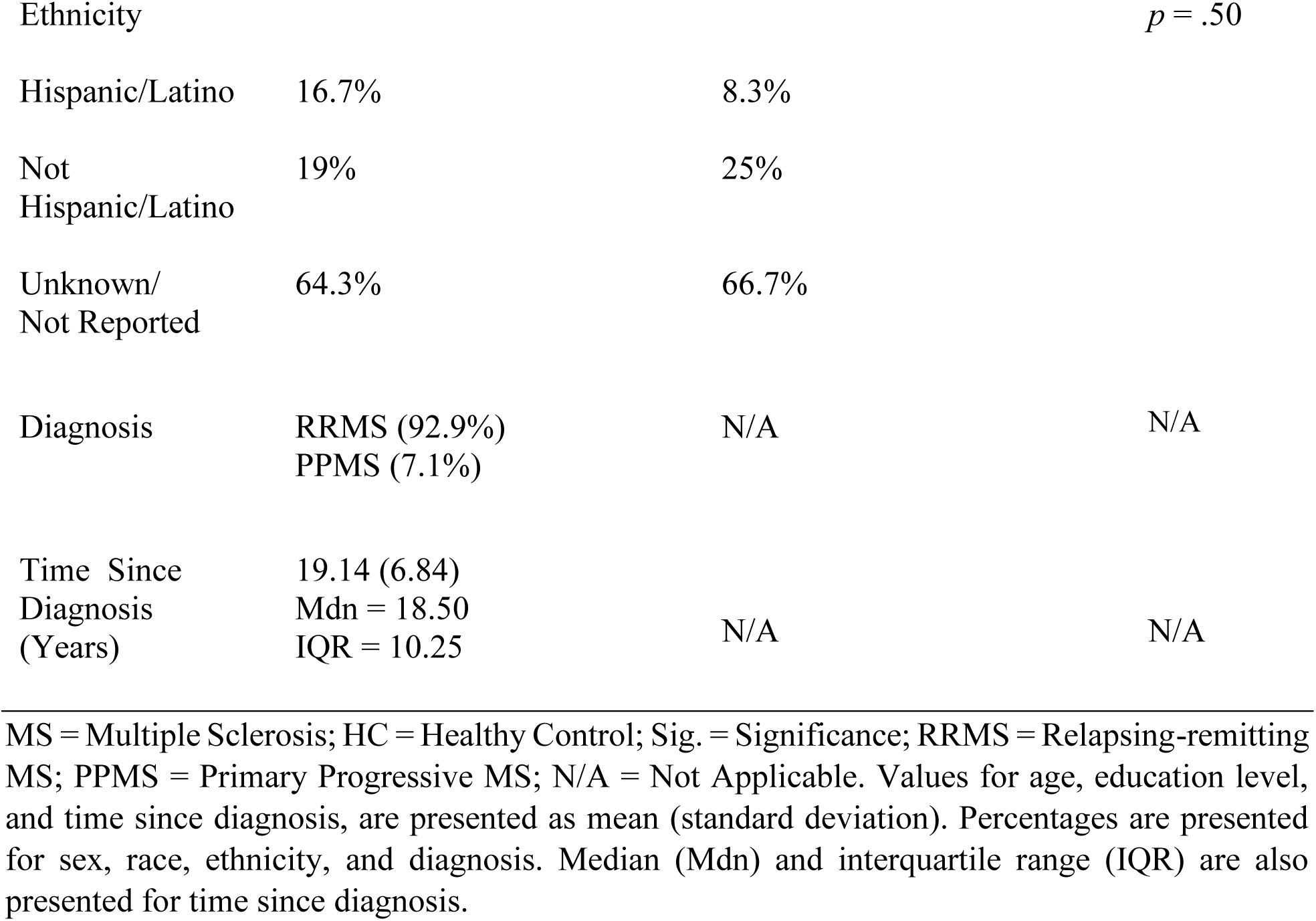
Study Sample Demographics.

### 2.2. Neuropsychological test data

As described in detail in Section 2.4.1., the Wechsler Adult Intelligence Scale, Fourth Edition (WAIS-IV) Digit Span test (Wechsler, 2008) and the Hopkins Verbal Learning Test – Revised (HVLT-R; Benedict et al., 1998) were used as component measures for generating a measure of WM performance.

Two subtests from the WAIS-IV Digit Span were administered: Digit Span Forward (DSF) and Digit Span Backward (DSB). During DSF, an examiner reads aloud a string of single-digit numbers to the participant who must then repeat back the digits in the same order. Across a total of 16 trials, the trials gradually increase in string length, beginning with two digits and ending with nine digits. A similar testing format occurs for the DSB subtest, except the participant must repeat the digits back in the reverse order from which they were read by the examiner. Both tests end when the participant incorrectly responds to two consecutive trials of the same string length. Thus, DSF assesses active storage of information, while DSB assesses manipulation of stored information.

The HVLT-R is a test of verbal learning and memory. An examiner reads aloud a list of 12 words that belong to three semantic categories. The participant must then immediately repeat as many words as they can remember in any order. The procedure is repeated across three learning trials followed by one delayed recall trial. Immediate recall of Trial 1 is equivalent to simple span WM tasks (Unsworth & Engle, 2006) and is included as a measure of immediate WM in the present investigation.

### 2.3. fMRI data acquisition

A 3T Siemens (Erlangen, Germany) MAGNETOM Skyra scanner was used for neuroimaging data acquisition in all parent studies. Magnetization Prepared Rapid Gradient Echo (MPRAGE) sequences (1 mm isotropic voxels; two parent studies: TR: 2100 ms; TE: 3.43 ms; one parent study: TR: 2100 ms; TE: 2.98 ms) were used to acquire T1-weighted anatomical images in each of the three studies. Thirty-two echo-planar, interleaved, functional slices were acquired with the following parameters for two of the parent studies: 3 mm isotropic voxels, interslice gap: 0.3 mm, TR: 2000 ms, TE: 30 ms, field of view: 220 mm, flip angle: 90°. For the remaining study, 64 echo-planar, interleaved, functional slices were acquired with the following parameters: 2.2 mm isotropic voxels, interslice gap: 0.8 mm, TR: 785 ms, TE: 37.2 ms, field of view: 224 mm, flip angle: 55°. Data acquisition for this study employed a multiband accelerated factor of 8 to increase the number of volumes acquired during a single scan session. Thus, 522 volumes were acquired during the single resting state run for participants in this study; whereas 180 volumes were acquired during the single resting state run for participants in the other two studies (which did not employ multiband during acquisition). Analysis methods for accounting for the differing number of volumes across participants are further explained in Section 2.4.2.

### 2.4. Data analysis

#### 2.4.1. Behavioral data analysis

SPSS statistical software (v. 29.0; IBM Corp., Armonk, NY) and RStudio (v. 4.2.3; R Foundation for Statistical Computing; Vienna, Austria) were used for data analysis. Independent-samples t tests examined group differences in age, years of education, and WM performance (see WM performance score below). Chi-square tests examined the same for biological sex, race, and ethnicity.

Principal component analysis (PCA) was conducted using raw scores from the three neuropsychological assessment scores operationalized as measures of WM (DSF, DSB and HVLT T1), resulting in a single WM component that served as our measure of WM capacity. These tests were used to represent the component processes of storage and manipulation within WM. DSF requires storage of increasingly longer sets of numbers; DSB requires manipulation of those sets into backward sequences; and HVLT potentially involves both storage and manipulation (if participants employ semantic clustering when repeating words back during recall). For meaningful interpretation, PCA scores were converted to z-scores using the HC mean and standard deviation (*M* = 0.46, *SD* = 0.91).

#### 2.4.2. fMRI data analysis

##### 2.4.2.1 Regions of interest

Given our *a priori* hypotheses regarding the roles of caudate and hippocampus RSFC in predicting WM performance, we generated four seeds – left and right sides for each region (Fig. 1). We used the right and left hippocampus and right and left caudate ROIs in the CONN Toolbox’s (RRID: SCR_009550; Whitfield-Gabrieli & Nieto-Castañón, 2012; Morfini, Whitfield-Gabrieli, & Nieto-Castañón, 2023) default atlases (the Harvard Oxford and AAL atlases). These four ROIs were used in subsequent higher-level analyses to generate connectivity maps between each seed and every voxel throughout the brain.

**Fig. 1.**
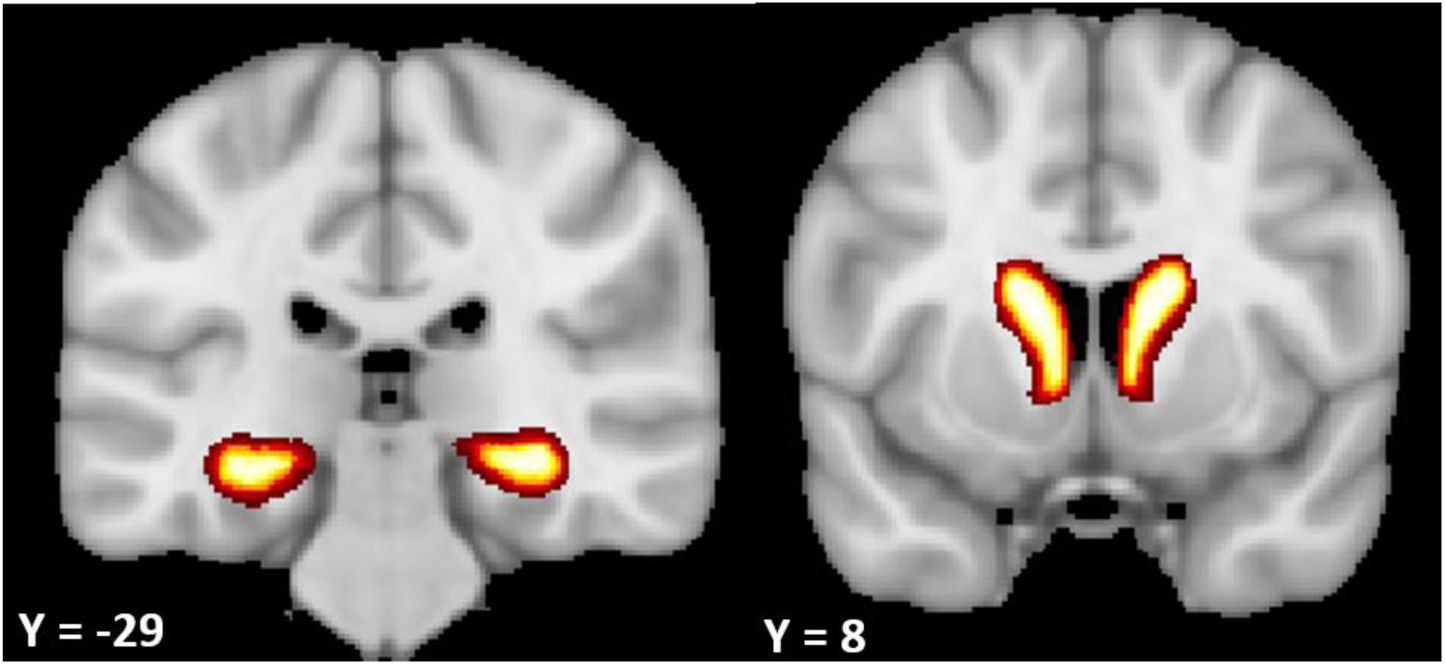
Bilateral hippocampus (left) and bilateral caudate nuclei (right) seeds used in all seed-to-voxel RSFC analyses. All seeds were derived from the Harvard Oxford and AAL atlases. Slice numbers correspond to MNI coordinates. Images are displayed in neurological orientation.

##### 2.4.2.2. Resting-state functional connectivity analysis

All preprocessing and resting-state analyses were performed using the CONN Toolbox. Functional (BOLD) data were co-registered to anatomical (T1) data before being normalized to MNI-152 space (1 mm). Segmentation involved parcellating the data into gray matter, white matter, and cerebrospinal fluid using CONN’s aCompCor tool. CONN’s Artifact Removal Tool (ART) was used to detect and remove volumes exhibiting large degrees of motion. Suprathreshold framewise displacement (FD) was used to detect such outlier volumes. Volumes displaying a global signal *z* value greater than 3 and FD across the entire scanning session greater than 0.3 mm were removed. We opted for these conservative motion parameters, given the inherent susceptibility of resting-state data to motion artifacts (Parkes et al., 2018) and due to the large size of our sample, which afforded us the ability to be more stringent. Data were then smoothed using a kernel of 5 mm FWHM. During denoising, first-order derivatives and quadratic effects of all motion regressors were included in the general linear model (GLM). Linear detrending was employed, and a bandpass filter of [0.008, 0.09] Hz was applied to the data to further control for influence from extraneous low and high frequency.

For first-level analysis, seed-to-voxel maps were generated for each of our four ROIs (bilaterial hippocampi and bilaterial caudate nuclei) for each participant. These were then entered into a group-level analysis, in which we included a group regressor (MS, HC) and two additional covariates. The first covariate accounted for the different acquisition timings within the constituent datasets (TR = 2000 ms in two studies and TR = 785 ms in one study). The second covariate accounted for resting state sequence order in the datasets – resting-state sequences were acquired prior to task sequences in two studies and after task sequences in one study. Linear contrasts examined differences in resting-state connectivity both between groups and within each group. For all analyses, a cluster-defining threshold of α = .001 and a cluster-correction threshold of α = .05 (familywise error) were used. The Harvard-Oxford and AAL atlases were used for cortical and subcortical neuroanatomical classification of functional activity, while the Spatially Unbiased Atlas Template of the Cerebellum (SUIT; Diedrichsen et al., 2009) was used for cerebellar classification.

To assess relationships between WM performance and RSFC with the caudate, hippocampus, and other brain regions, we conducted Pearson correlation analyses between the parameter estimates of each significant connection and WM performance score. We then corrected for the number of analyses using the Benjamini-Hochberg method to control the False Discovery Rate (α_FDR_ = .05). To investigate whether these connections predicted WM performance, we entered each surviving significant connection into a hierarchical multiple regression. In each of these models, WM performance score was the outcome variable. The first level adjusted for the demographic influences of age (centered), sex, and years of education (centered) on WM performance. The next level assessed the significance of the functional connection of interest after adjusting for these demographic influences.

## 3. Results

### 3.1. Group characteristics

As displayed in Table 1, there were no significant group differences in age [*t*(76) = 1.55. *p* = .12, *d* = 0.35], years of education [*t*(76) = -0.10. *p* = .92, *d* = 0.02], biological sex [*X^2^*(1) = 0.40. *p* = .53], race [*X^2^*(4) = 8.84, *p* = .07], or ethnicity [*X^2^*(2) = 1.38, *p* = .50] in the entire sample (N = 78). Our MS sample comprised predominantly of individuals with the relapsing-remitting phenotype (92.9%) and with an average time course of 19.14 years with the disease (*SD* = 6.84, Mdn. = 18.50, IQR = 10.25). Similar results emerged for the subset of participants with WM data available (N = 41). For this subset, there were no group differences in age [*t*(39) = 1.91. *p* = .07, *d* = 0.72], education [*t*(39) = -1.03. *p* = .31, *d* = 0.33], biological sex [*X^2^*(1) = 0.004. *p* = .95], race [*X^2^*(4) = 6.60. *p* = .16], or ethnicity [*X^2^*(2) = 2.21. *p* = .33]. 88.5% of this subset’s MS sample comprised of individuals with RRMS and with an average time course of 18.92 years (*SD* = 5.73, Mdn. = 19.00, IQR = 9.50).

### 3.2. MS participants displayed worse working memory performance

Compared to HC participants (*M* = 0.00, *SD* = 1.00), MS participants (*M* = -0.79, *SD* = 1.06) demonstrated worse WM performance, as indicated by significantly lower WM performance scores: *t*(39) = 2.37, *p* = .02, *d* = 0.76 (Fig. 2).

**Fig. 2.**
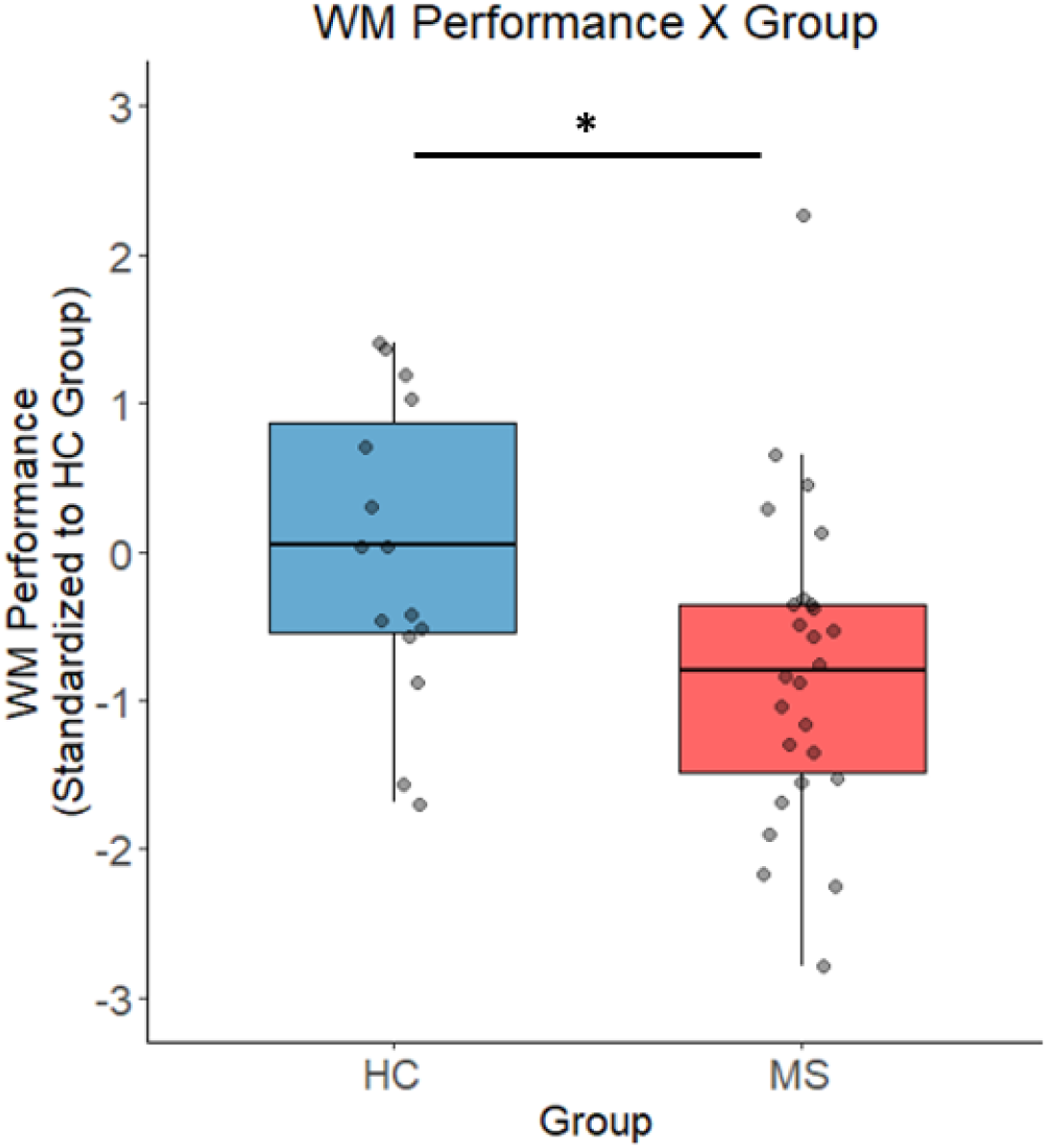
Working memory performance between groups, as measured by a performance score standardized to the mean of the HC group. MS participants (red) displayed significantly lower scores than HC participants (blue). Points represent individual scores. MS = Multiple Sclerosis; HC = Healthy Control; WM = working memory. N = 41 (26 MS, 15 HC). \**p* < .05

### 3.3. Within-group and between-group resting-state functional connections for each hippocampus and caudate seed

Brain regions displaying significant functional connectivity at rest with each seed region are displayed in Table 2. The sole group difference emerged in the right hippocampus. MS participants displayed significantly stronger connectivity between the right hippocampus and left dorsomedial prefrontal cortex (dmPFC), compared to HC participants (Fig. 3A).

**Fig. 3.**
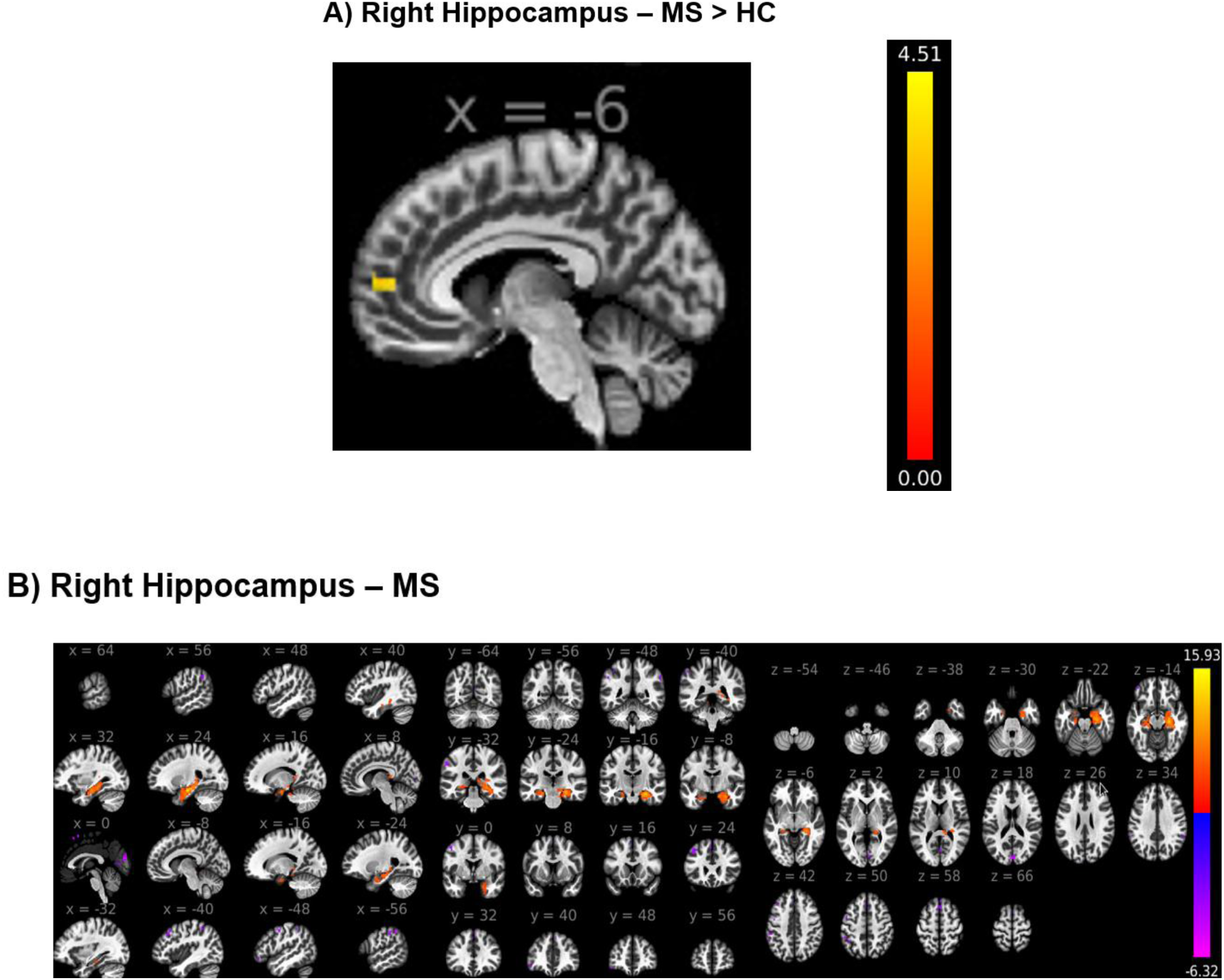
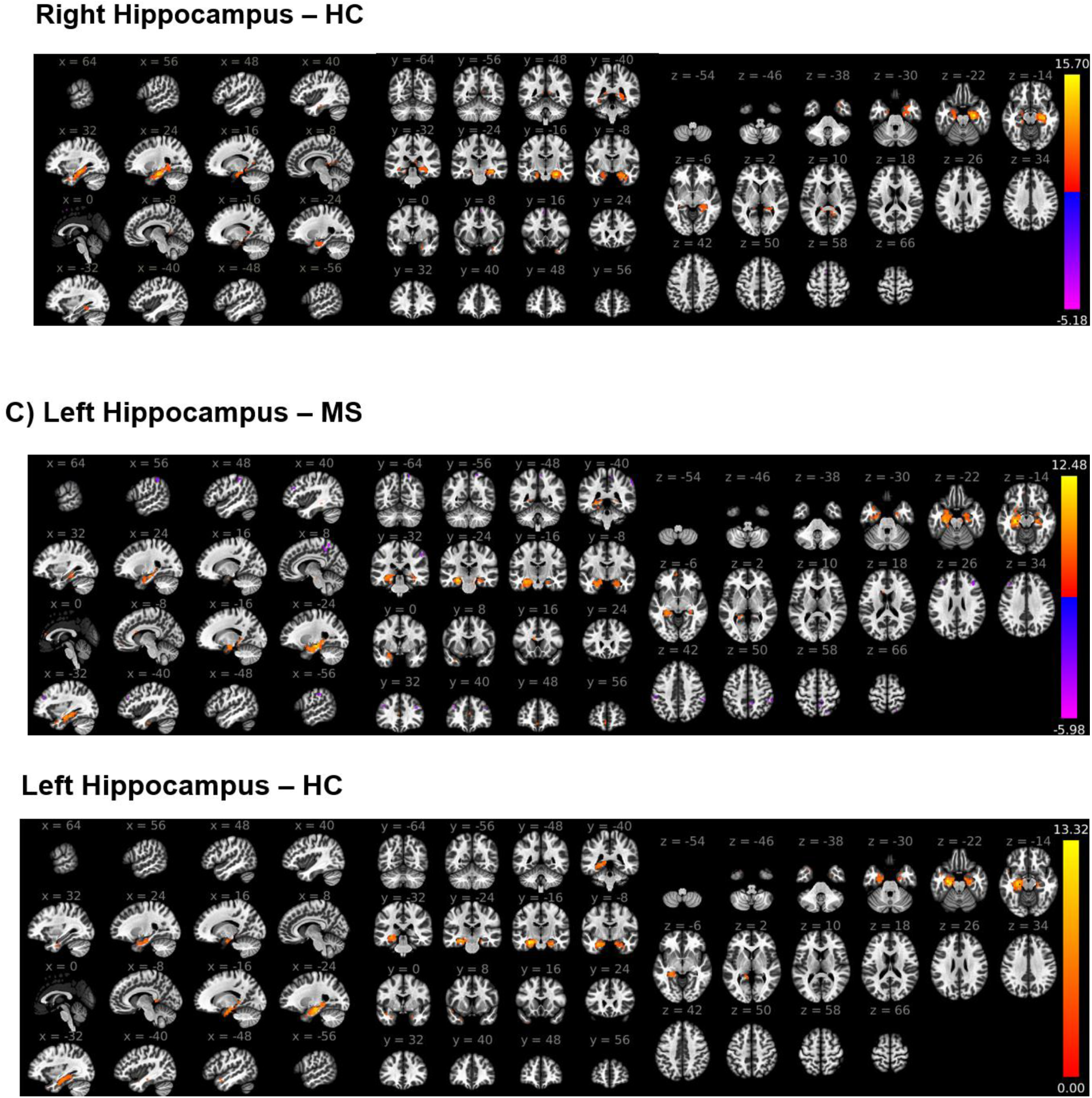
Whole-brain voxelwise RSFC results with hippocampus seeds. A) MS participants displayed stronger functional coupling between the right hippocampus and left dmPFC at rest, compared to HC participants. Within-group results for each group are also displayed for the right hippocampus (Panel B) and left hippocampus (Panel C). T-statistic maps are displayed [cluster-defining threshold: α = .001, cluster-correction threshold: α_FWE_ = .05). Warmer colors indicate regions sharing connectivity with the respective hippocampal seed, and cooler colors indicate dysconnectivity. Slice numbers correspond to MNI coordinates. Images are displayed in neurological orientation.

**Table 2.**
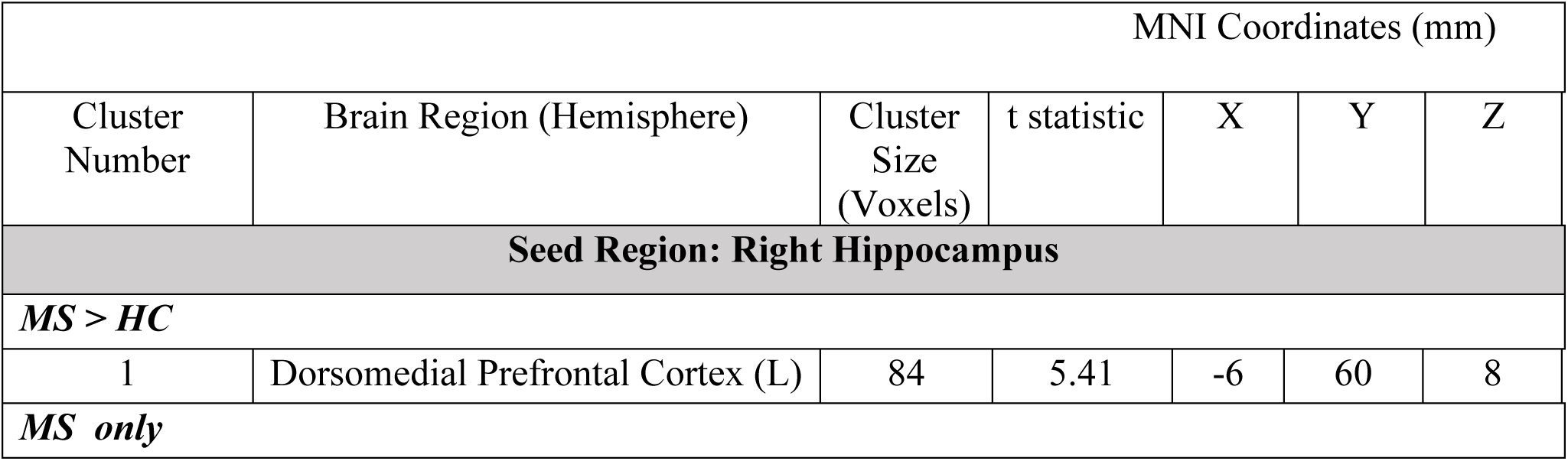
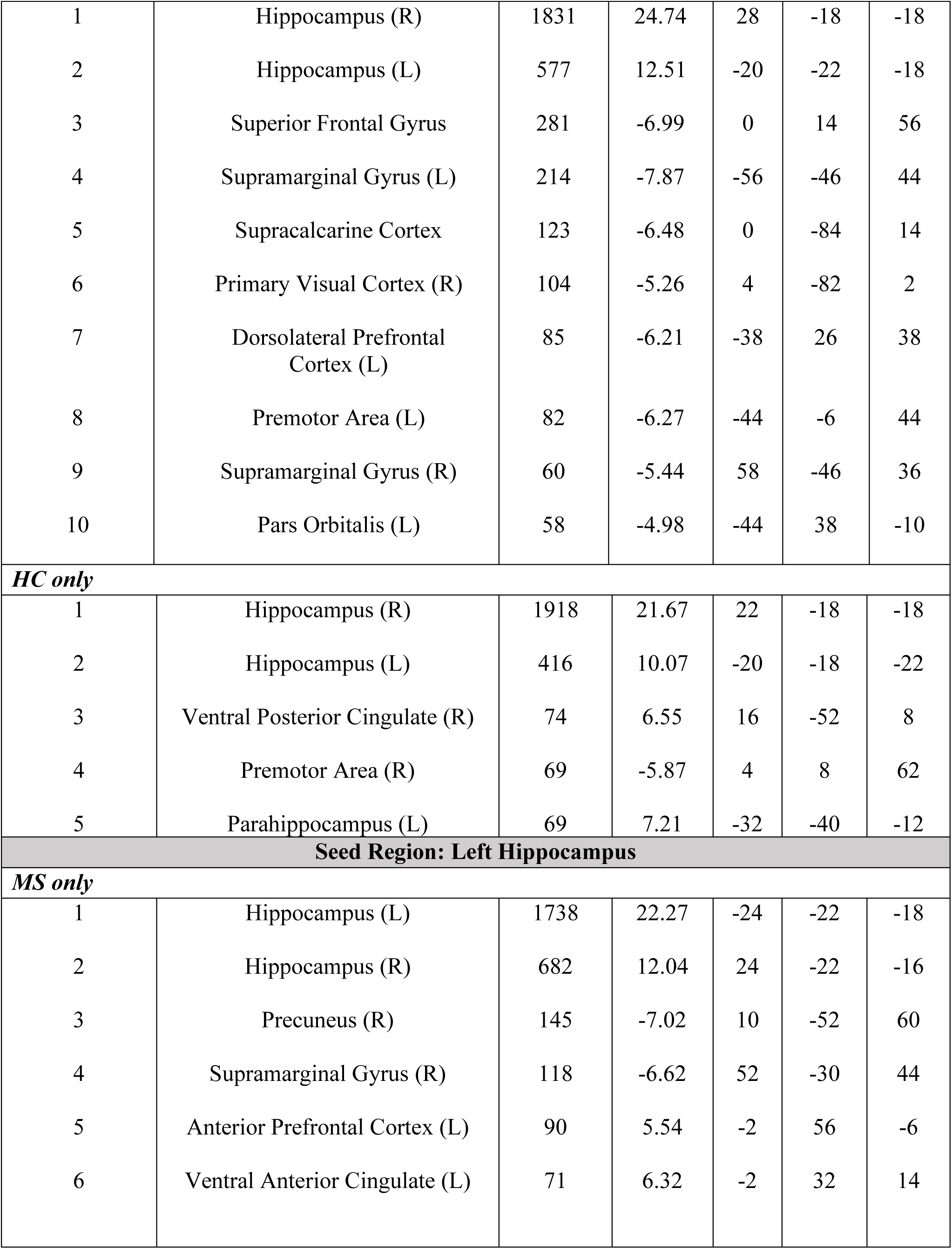
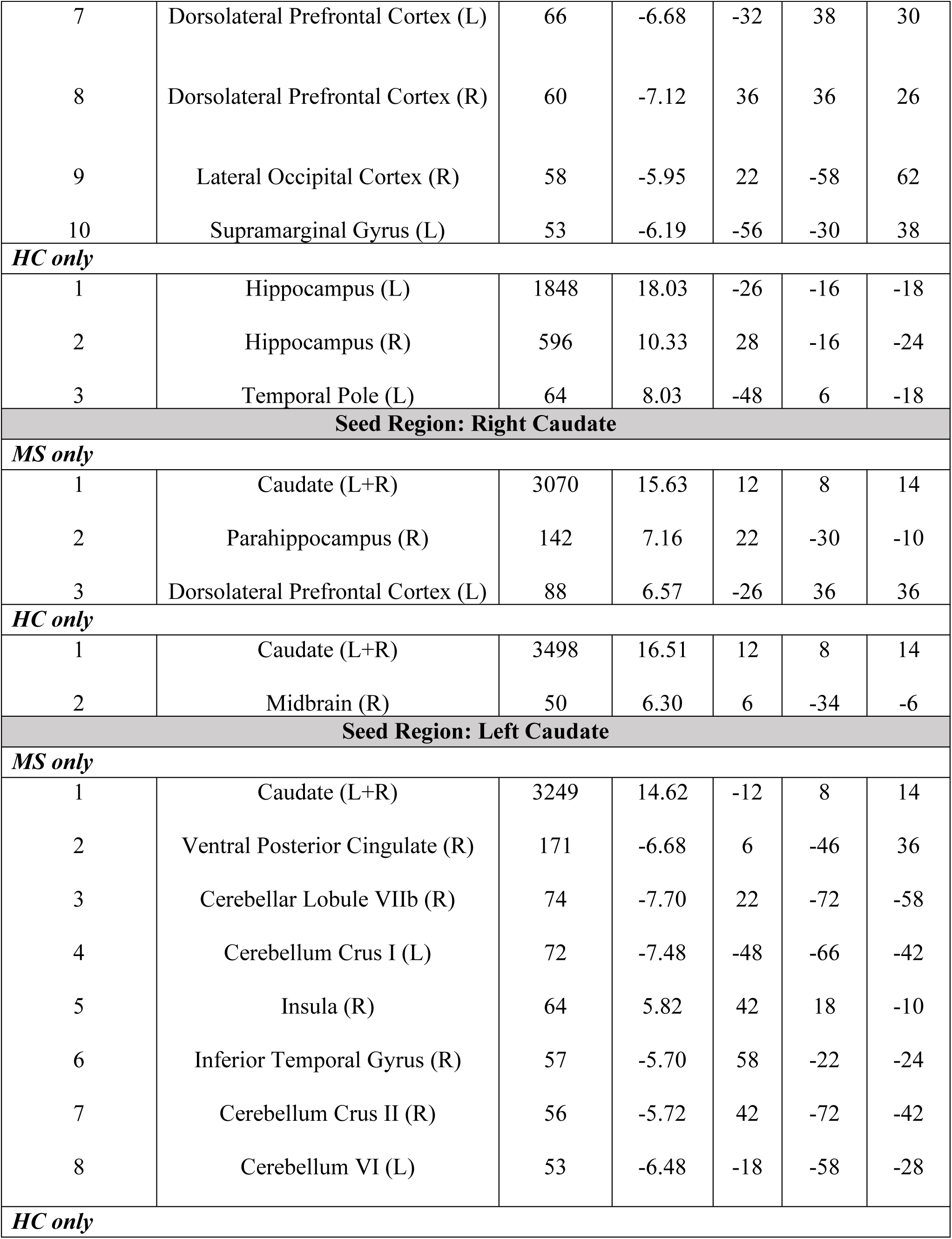
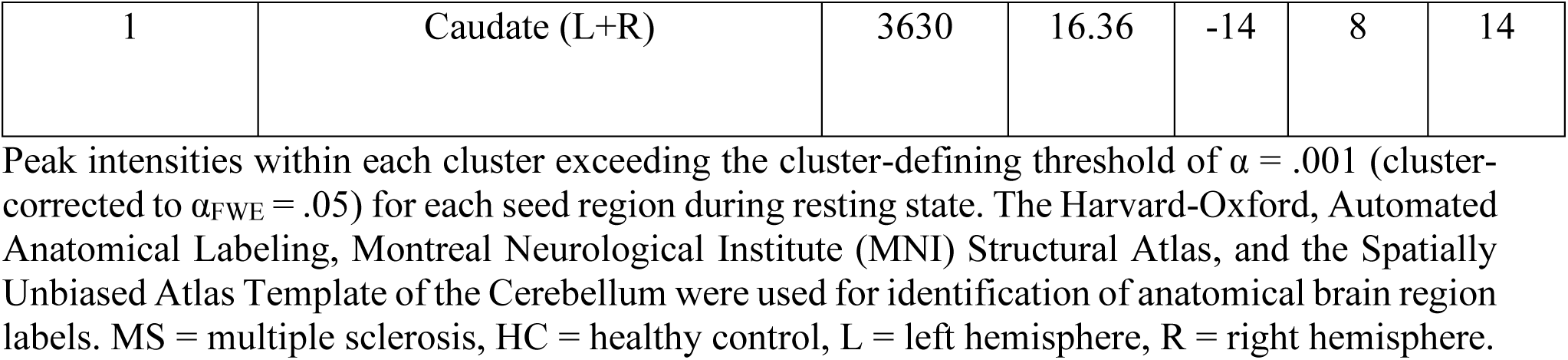
Resting-state functional connectivity analysis clusters and local maxima (i.e., peak) coordinates for each hippocampus and caudate seed region.

With respect to within-group differences, MS participants displayed dysconnectivity between the right hippocampus and the superior frontal gyrus, left supramarginal gyrus (SMG), supracalcarine cortex, right primary visual cortex, left dorsolateral prefrontal cortex (dlPFC), left premotor area (PMA), right supramarginal gyrus (SMG), and left pars orbitalis (Fig. 3B). HC participants displayed connectivity between the right hippocampus and the right ventral posterior cingulate (vPCC), and left parahippocampus, as well as dysconnectivity with the right PMA. For the left hippocampus (Fig. 3C), MS participants displayed connectivity with the left anterior PFC, and left ventral anterior cingulate (vACC), as well as dysconnectivity with the right precuneus, right SMG, left dlPFC, right dlPFC, right lateral occipital cortex, and left SMG. HC participants displayed only connectivity between the left hippocampus and the left temporal pole.

Both groups displayed sparse connectivity of the caudate with the rest of the brain. Within the MS group, connectivity emerged between the right caudate and the right parahippocampus and left dlPFC (Fig. 4A). HC participants only displayed connectivity with a small region of the brain stem. For the left caudate seed (Fig. 4B), MS participants displayed connectivity with the right insula, as well as dysconnectivity with the right vPCC and right inferior temporal gyrus (ITG). They also displayed dysconnectivity with a number of cerebellar subregions, including the right Lobule VIIb, left Crus I, right Crus II, and left IV region. HC participants only displayed connectivity between the left caudate and the bilateral caudate nuclei.

**Fig. 4.**
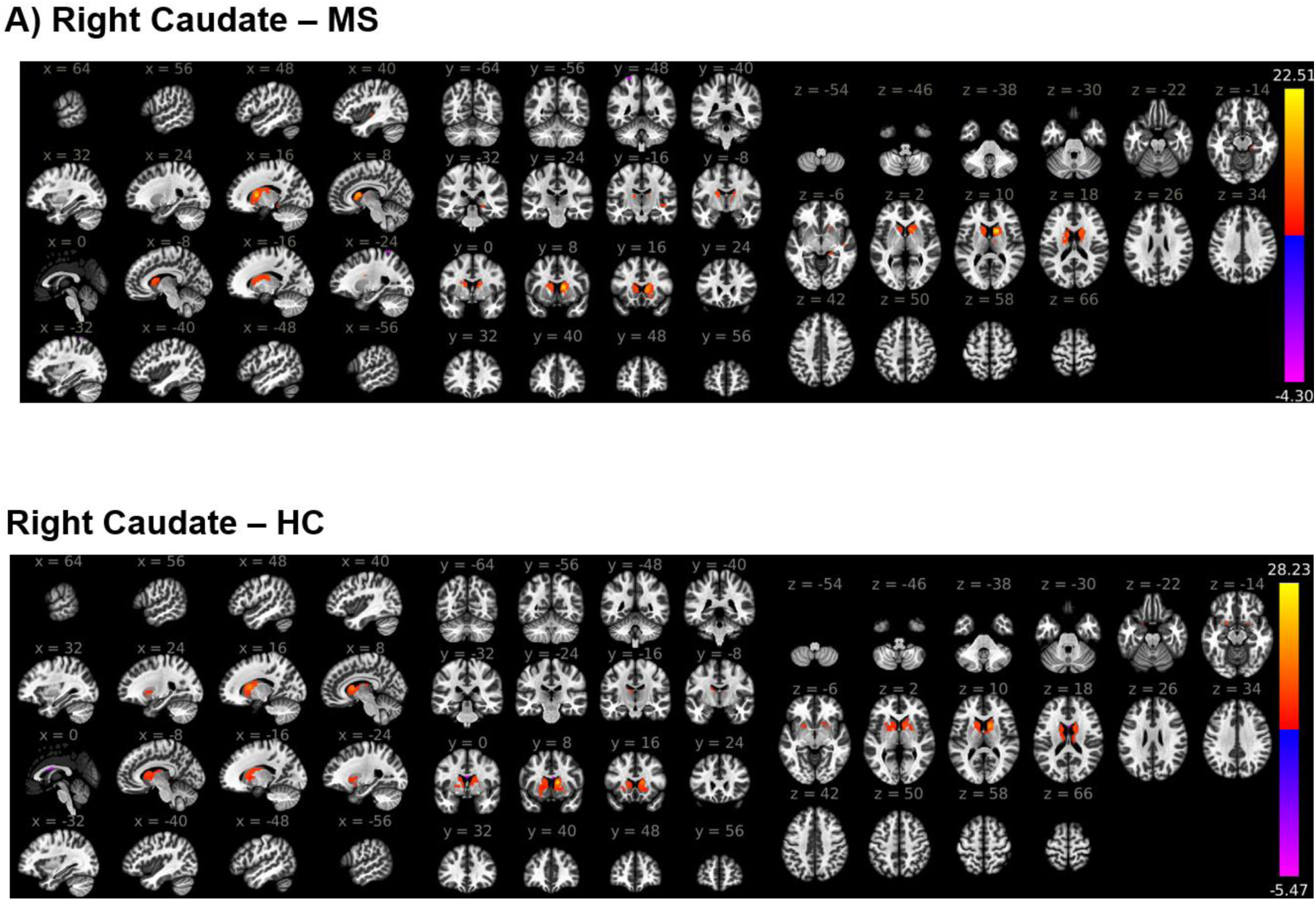
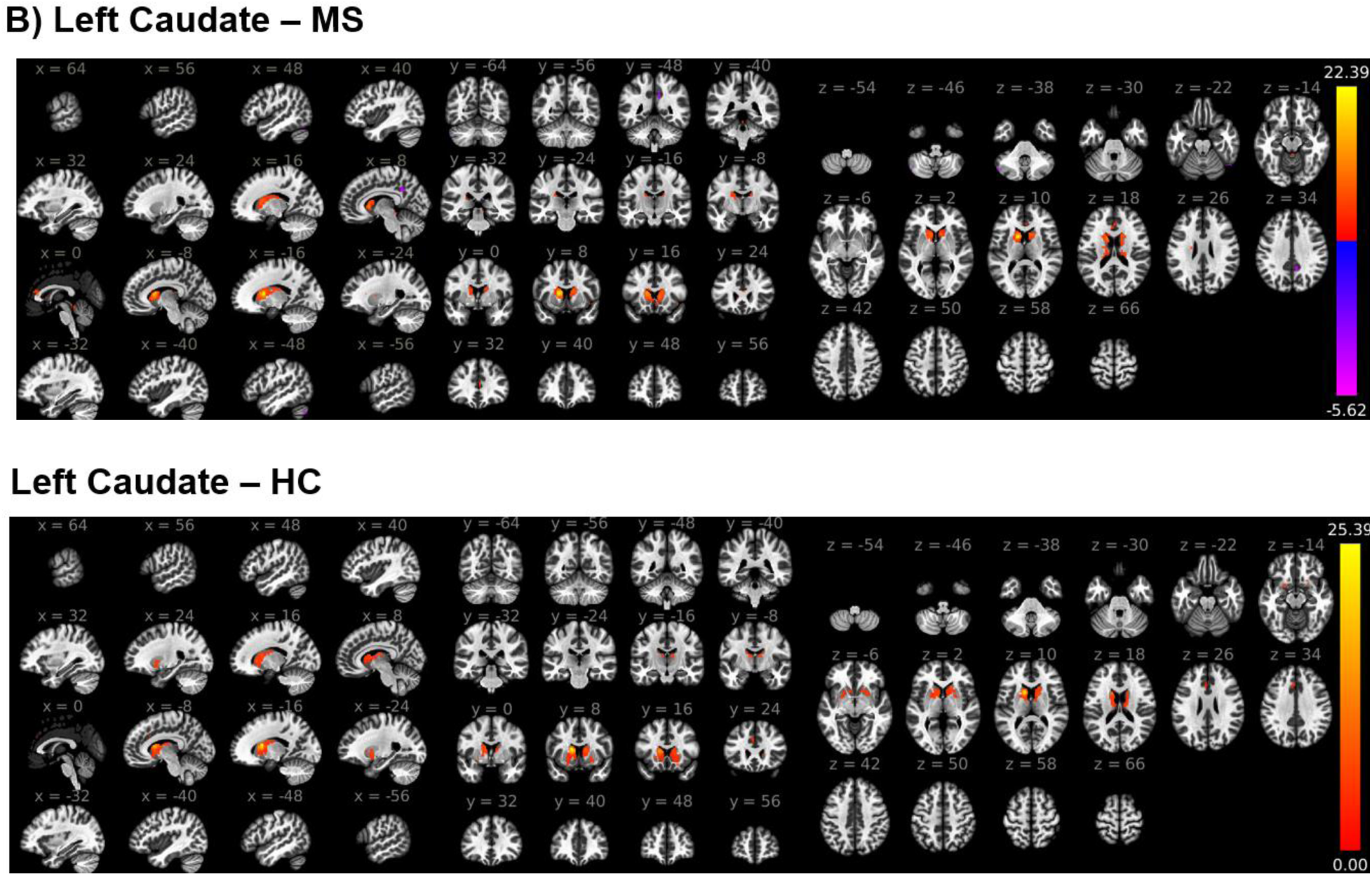
Whole-brain voxelwise RSFC results with caudate seeds. Within-group results for each group are displayed for the right caudate (Panel A) and left caudate (Panel B). T-statistic maps are displayed [cluster-defining threshold: α = .001, cluster-correction threshold: α_FWE_ = .05). Warmer colors indicate regions sharing connectivity with the respective caudate seed, and cooler colors indicate dysconnectivity. Slice numbers correspond to MNI coordinates. Images are displayed in neurological orientation.

### 3.4. MS participants display hippocampus and caudate resting-state functional connections that predict WM performance

Results of all correlation analyses testing the association between the above functional connections and WM performance in each group are displayed in the Supplementary Material. The significantly greater RSFC between the right hippocampus and left dmPFC in MS participants, compared to HC participants, did not correlate with MS participants’ significantly worse WM performance [*r*(39) = -.10, *p*_uncorr_ = .55]. Within the MS group (Fig. 5A), stronger connectivity between the left hippocampus and left vACC was associated with lower WM performance scores [*r*(24) = -.49, *p*_uncorr_ = .01, *p*_corr_ = .054]. Meanwhile, stronger connectivity between the left hippocampus and left SMG [*r*(24) = .49, *p*_uncorr_ = .01, *p*_corr_ = .054] and between the left caudate and right insula [*r*(24) = .48, *p*_uncorr_ = .01, *p*_corr_ = .10] were each associated with better WM performance scores. As a follow-up analysis (Fig. 5B), we also tested whether these three connections were significantly related to each other. This was the case with the hippocampal connections, which exhibited a moderate association [LHipp/LvACC and LHipp/LSMG: *r*(40) = -.45, *p* = .003]. The left caudate-right insula connection, however, did not correlate with either hippocampal connection [LCaud/RInsula and LHipp/LvACC: *r*(40) = .24, *p* = .12; LCaud/RInsula and LHipp/LSMG: *r*(40) = -.20, *p* = .21]. No correlations with WM performance emerged for the right hippocampus and right caudate in the MS sample.

**Figure 5.**
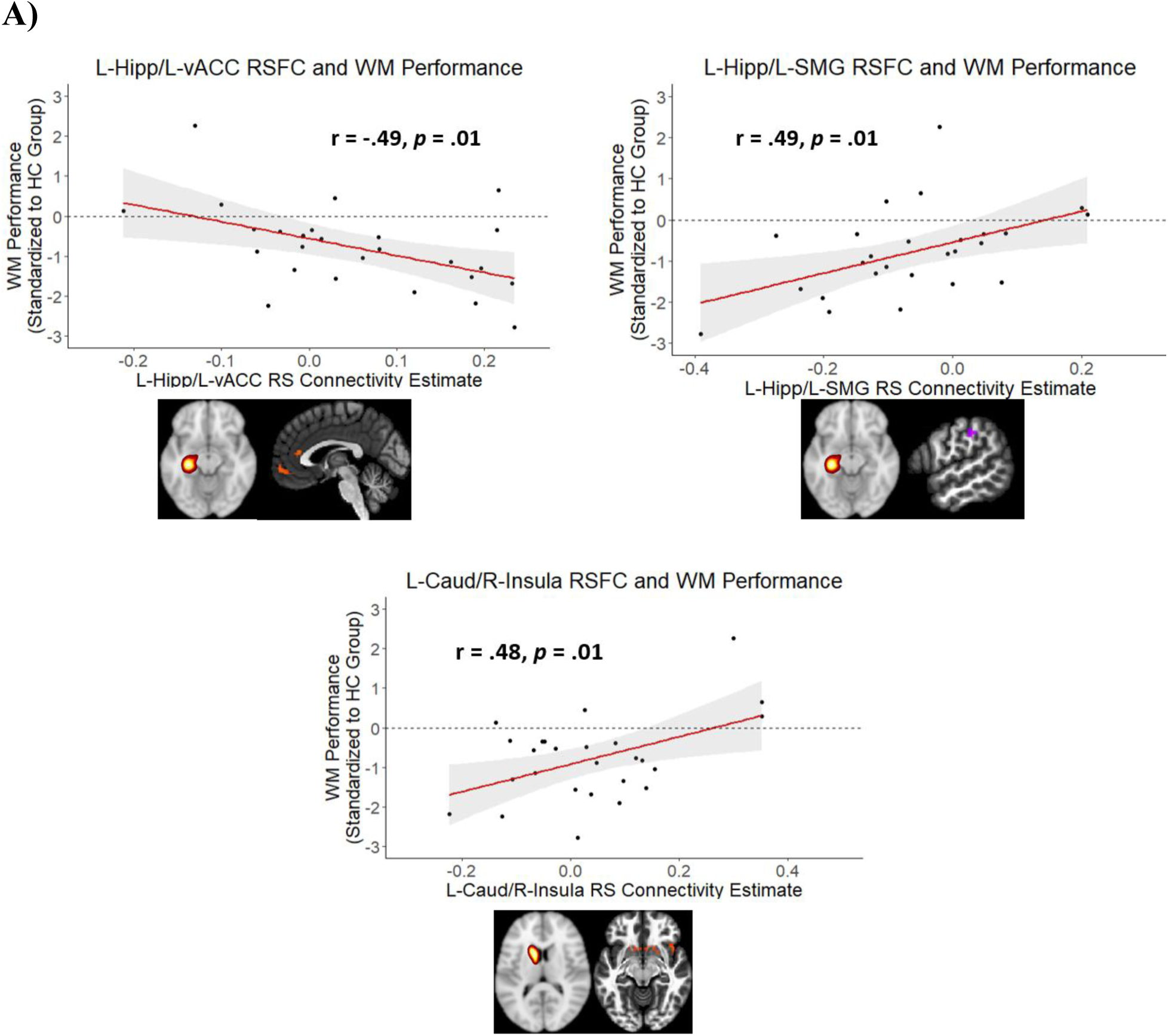
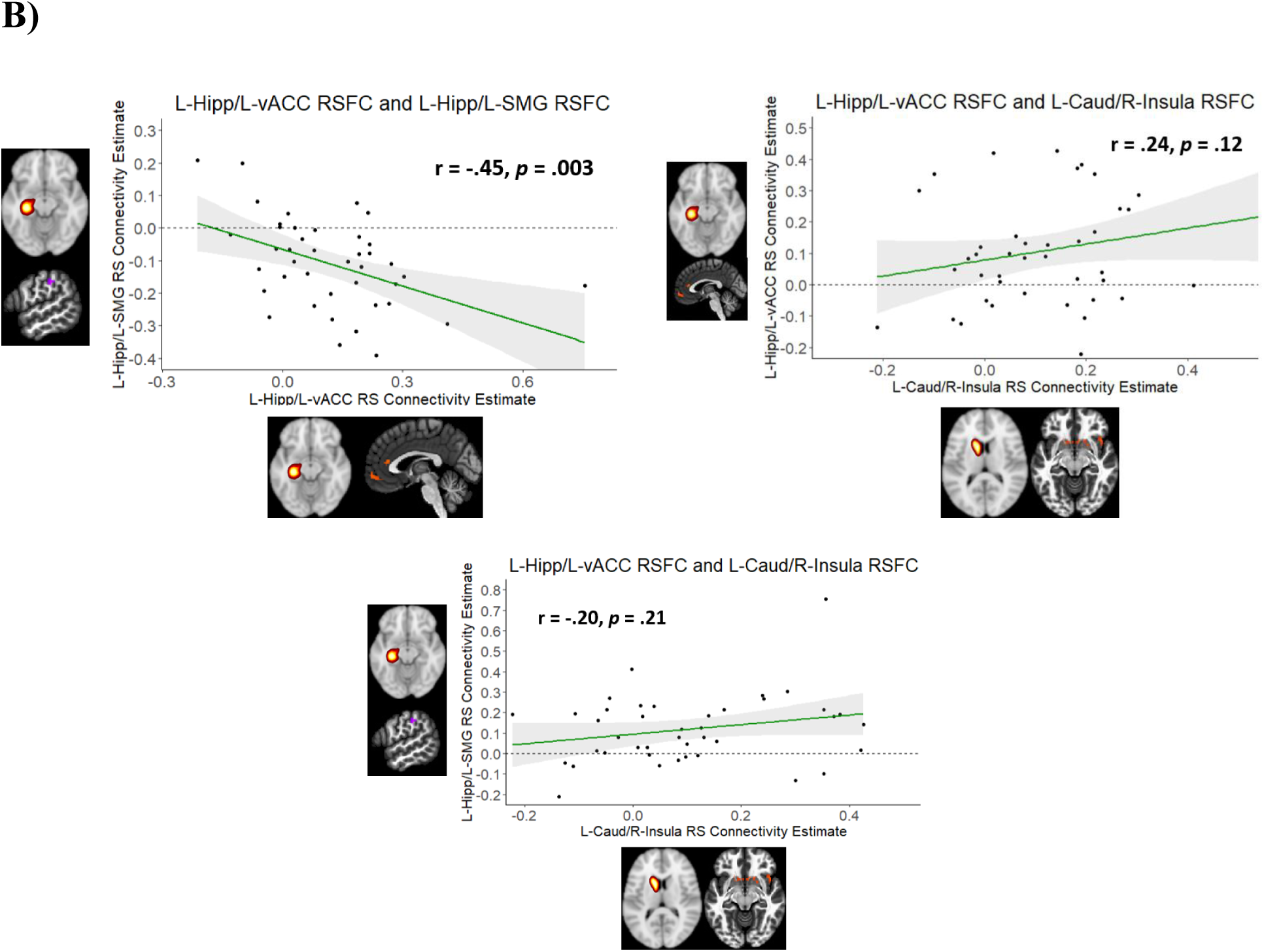
A) Significant hippocampus and caudate functional connections at rest associated with working memory performance in MS participants. Stronger RSFC between the left hippocampus and left vACC was associated with worse WM performance (i.e., lower scores) in MS participants. However, stronger RSFC between the left hippocampus and left SMG and between the left caudate and right insula were each associated with better WM performance (higher scores) in MS participants. B) Furthermore, connectivity between each of the hippocampal connections was associated with each other, while neither hippocampal connection correlated with the caudate connection. *Note:* Panel A analysis n = 26; Panel B analysis n = 42. Different sample sizes are due to limiting factor of neuropsychological test data availability for the three parent datasets used in the current study. WM = working memory; RSFC = resting state functional connectivity; L-Hipp = left hippocampus; L-vACC = left ventral anterior cingulate cortex; L-SMG = left supramarginal gyrus; L-Caud = left caudate.

In HC participants, there was no association between any resting-state functional connection (with any seed region) and WM performance. No correlations with WM performance score emerged for the left hippocampus, right hippocampus, left caudate, or right caudate in this group.

We then conducted follow-up hierarchical regression models to test whether each resting state functional connection indeed predicted WM performance. For the MS sample, despite the corrected *p* value for each of the three connections exceeding the FDR threshold of .05, we opted to submit them for regression analysis. Our rationale for this approach was due to the moderate effect sizes (*r*’s = -.49, .48, .49) of the relationship that each shared with WM performance. Thus, we opted to include them to minimize the effects of a Type II Error unknowingly made during our FDR correction procedures during the correlation analyses.

Regression results indicated that these resting state connections did predict WM performance in MS participants. Connectivity between the left hippocampus and left vACC at rest accounted for an additional 24% of variance in WM performance scores above and beyond the influences of age, sex, and education level. Stronger connectivity between these regions at rest predicted worse WM performance scores (β = -0.50, *t* = -2.67, *p* = .01). Additionally, connectivity between the left hippocampus and left SMG at rest accounted for an additional 32% of the variance in WM performance scores above and beyond demographic factors. However, stronger connectivity between these regions at rest predicted *better* WM performance (β = 0.64, *t* = 3.25, *p* = .004) in MS participants. Finally, connectivity between the left caudate and the right insula accounted for an additional 21% of variance in WM performance scores and also predicted better WM performance (β = 0.47, *t* = 2.37, *p* = .03). Results for each model are displayed in Table 3.

**Table 3.**
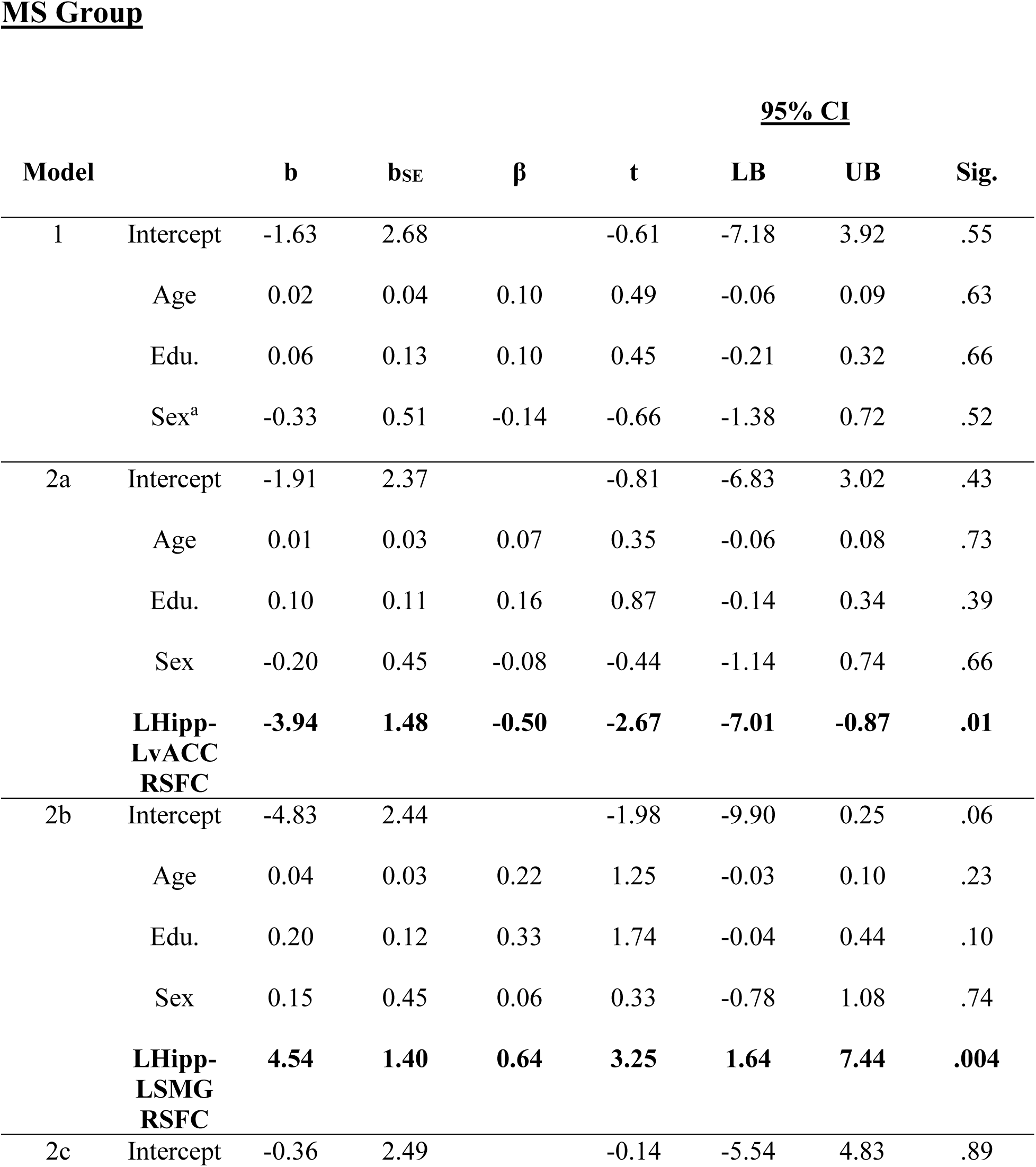
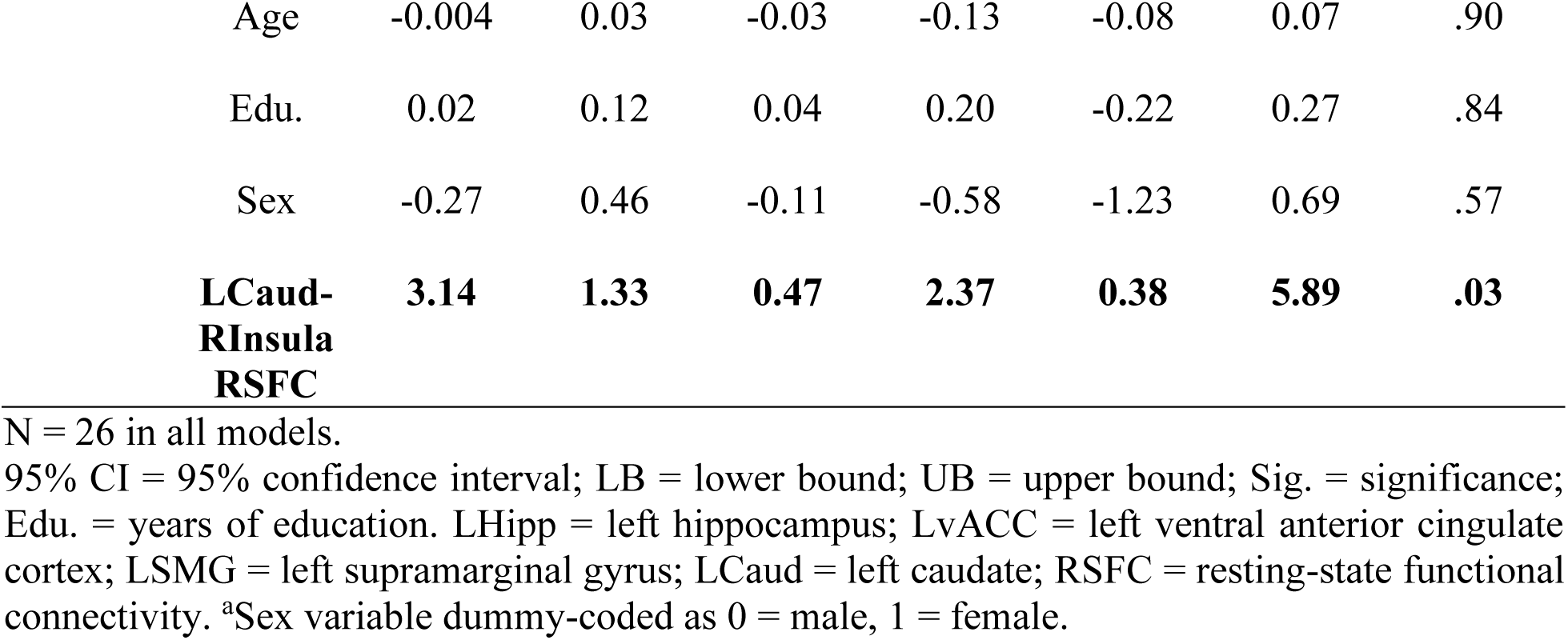
Regression model results for each significant resting-state functional connection in predicting working memory performance, as measured by WM performance score (outcome variable in all models). Model 1 includes only demographic covariate regressors, while each Model 2 (2a, 2b, 2c) tests the independent contribution of each resting-state functional connection identified during correlation analysis.

## Discussion

The present study tested whether seed-based, voxelwise functional connectivity with the hippocampus and caudate at rest predicts WM performance in people with MS. These two regions were selected, given their mutual contributions to WM (Freedberg, 2023; Müller et al., 2018). Our results support this hypothesis. MS participants displayed worse WM performance, compared to HC participants, as reflected by significantly lower scores on the WM performance measure. However, stronger connectivity between the right hippocampus and left dmPFC that was also observed in MS participants did not contribute to this difference in WM performance. Within MS participants, three connections – two with the left hippocampus and one with the left caudate – emerged whose activity at rest each predicted WM performance. Connectivity between the left hippocampus and left SMG and between the left caudate and right insula each predicted better WM performance, while that between the left hippocampus and left vACC predicted worse performance. Furthermore, connectivity between both hippocampus connections correlated with each other. Collectively, these results not only identify potential hippocampus and caudate circuits whose intrinsic connectivity can predict WM performance in MS, but they also identify a potential resting-state network centered around the left hippocampus that may serve as an important neural marker to monitor for the development of WM impairment throughout the course of MS progression.

### 4.1. Hippocampal and caudal connections at rest predict WM performance in MS

Our results indicate that resting-state functional connections involving the hippocampus and the caudate predict WM performance in MS, which partially aligns with evidence of interactions between the two regions during working memory tasks in healthy individuals (Freedberg, 2023; Müller et al., 2018). In the present study, however, RSFC between the hippocampus and caudate was not prominent. Increased functional coupling between the right caudate and right parahippocampus was the only connection to emerge, but this connectivity was not associated with WM performance. This is likely due to this connection occurring at rest, rather than being directly elicited by a WM task (as in task-based connectivity). It is possible that an episodic memory or WM task would elicit shared functional connections among the caudate and hippocampal circuits identified in the current study, mirroring previous interactions between them during episodic memory reported elsewhere (Freedberg et al., 2022; Freedberg, 2023; Müller et al., 2018). The left caudate’s connectivity with the right insula at rest, however, did predict WM performance. The insula shares functional connections with the basal ganglia (Ghaziri et al., 2018), and it has been shown to be involved in WM (Kurth et al., 2010; Menon & Uddin, 2010), with some studies demonstrating its involvement in verbal WM specifically (Llorens et al., 2023). These studies, however, were primarily in neurologically healthy individuals. To our knowledge, our study is the first to identify *resting-state* insular activation as a potential neural marker of WM in MS.

L-caudate/R-insula connectivity at rest did not correlate with either L-hippocampus/L-vACC or L-hippocampus/L-SMG connectivity at rest. Tractography and RSFC studies indicate that the insula shares structural (Ghaziri et al., 2018) and functional (Ghaziri et al., 2024) connectivity with both the hippocampus and the caudate. While our findings do not align with this work, they do still indicate that caudate-insula RSFC may serve as a neural marker for WM performance in MS, albeit a distinct one that operates in *parallel* with the aforementioned hippocampal connections.

### 4.2. A potential resting-state hippocampus network for WM in MS

Perhaps unsurprisingly, the majority of our findings center specifically around RSFC of the left hippocampus. Verbal working memory is often associated with left lateralized brain activity (though a recent meta-analysis suggests otherwise (Emch, von Bastian, & Koch, 2019)). The three neuropsychological measures used to generate our WM performance measure – DSF, DSB, and Trial 1 of the HVLT – are also verbal in nature. Functional coupling between the left hippocampus and left vACC predicted worse WM performance, while that between the left hippocampus and left SMG predicted better WM performance. These three regions – the hippocampus, vACC, and SMG – have each been shown to underlie different components of the prominent multicomponent model of working memory (Baddeley & Hitch, 1974). In brief, this model posits that WM is a systematic integration of component and executive processes. Lower-level systems – the phonological loop and visuospatial sketchpad – encode and temporarily store verbal/auditory and visuospatial information, respectively. The activity between these two systems is coordinated by an attentional control system known as the central executive. An additional component, the episodic buffer, then integrates information collected from the various stores into distinct (but temporary), coherent episodes (Baddeley, 2000). Both the ACC and SMG have been suggested to contribute to the component processes of the multicomponent model. Reports have indicated a more distributed role of the ACC in underlying processes that resemble the central executive (Duma et al., 2019; Kaneda & Osaka, 2008; Lenartowicz & McIntosh, 2005), while the SMG appears to contribute primarily to processes underlying the phonological loop (Deschamps, Baum, Gracco, 2014; Purcell, Rapp, & Martin, 2021). Thus, the contributions of these two hippocampal connections’ RSFC to WM performance in MS observed in the current study are consistent with other task-based empirical studies for the multicomponent model of WM in healthy individuals. Furthermore, the ACC and the SMG also display intrinsic functional connectivity (i.e., at rest) with each other (Du et al., 2020), which aligns with the association between their respective hippocampal connections observed in the current study.

Our findings are also consistent with other reports of the role of the hippocampus in memory performance in MS. As in the current study, Hulst and colleagues (2015) demonstrated that stronger RSFC in the left hippocampus was a predictor of memory performance in people with MS. However, in that study, a memory composite measure was computed using tests that evaluated various forms of memory, including delayed or long-term memory components; whereas, in the current study, we attempted to isolate RSFC contributions of the hippocampus and caudate to WM specifically (no delayed or long-term memory components). Our left hippocampus/left vACC finding is also consistent with another study by the same group (Hulst et al., 2012) that demonstrated increased *local* activation within each of these regions, compared to healthy individuals, during the encoding phase of an episodic memory task in cognitively preserved people with MS. The authors interpret this activation as adaptive functional reorganization to offset cognitive decline caused by disease-related structural damage (Hulst et al., 2012). Eventually, after significant structural damage accumulates over time, this increased functional reorganization is no longer sustainable and leads to a “network collapse,” characterized by inefficient neural signaling and cognitive impairment (Hulst et al., 2012; Rocca et al., 2012; Schoonheim, Geurts, & Barkhof, 2010; Schoonheim, Meijer, & Geurts, 2015). Our results align with this explanation, as stronger connectivity between the hippocampus and vACC *at rest* was predictive of *worse* WM performance. Thus, the robust RSFC between the left hippocampus and left vACC revealed here might be one such neural marker that may signal an impending network collapse that may impact WM in people with MS.

The above interpretation also coincides with our left hippocampus/left SMG result. Greater coupling between these regions at rest predicted *better* WM performance in our MS sample. The left SMG is a focal region for the processing of verbal information (Deschamps et al., 2014; Purcell, Rapp, & Martin, 2021). Since our WM performance score was derived from verbal WM measures, it is therefore possible that connectivity between the left hippocampus and left SMG acts as the “primary” circuit for facilitating verbal WM in MS. This would explain the positive relationship between this circuit’s activity at rest and better WM performance. When disease pathology progresses to a point where this primary circuit cannot sustain WM performance, functional reorganization occurs in the recruitment of an alternative hippocampal circuit (with the left vACC). Because this secondary circuit may maintain performance, but at a suboptimal level of efficiency, its connectivity is predictive of worse WM performance (and serves as a neural marker for impending WM network collapse). This interpretation is supported by functional connections that the vACC and SMG also share with each other at rest in healthy individuals (Du et al., 2020).

The only group difference to emerge in RSFC was between the right hippocampus and left dmPFC, where MS participants displayed enhanced connectivity, compared to HC participants. However, this connectivity did not predict WM performance. The hippocampus and dmPFC are part of the default-mode network (DMN), a functionally connected set of regions whose activity is suppressed when we are actively engaged in a task but is otherwise online. Some studies have linked DMN aberrations to cognitive impairment in MS (Eijlers et al., 2017; Rocca et al., 2010, 2012). Furthermore, with respect to WM specifically, the dmPFC has been shown to encode emotional content for subsequent WM maintenance (Smith et al., 2018). Therefore, our findings may indicate an additional secondary circuit to test in a prospectively designed future study that can detect the contributions of right hippocampus/left dmPFC RSFC on WM performance in MS.

### 4.3. Limitations and future directions

The current study has some limitations that should be taken into account. First, this study was a secondary analysis of an aggregate dataset of three independent studies. Although we employed necessary controls during data analysis to account for inter-study sources of variability (e.g., a different TR in one parent study), it is possible that this variance could have still influenced findings. As mentioned previously, future prospectively designed studies will likely further characterize the connections we have presented here as well as possibly identify additional ones. Second, as an additional limitation of a secondary analysis, our brain-behavior analyses were constrained by the number of participants with available neuropsychological test data to compute our WM performance score. As a result of the smaller samples available for those analyses, it is possible we lacked sufficient statistical power to identify additional RSFC patterns predictive of WM performance. Finally, the current study’s findings are limited to verbal working memory, specifically. It is possible that part of our results – specifically, the relationship between left hippocampus/left SMG RSFC and WM performance – are a result of the domain under investigation. Future work should focus on determining whether the results reported here are domain-general (across all WM types) or are specific to the domain of verbal WM. Furthermore, if the latter, future work should also determine whether verbal WM performance measures derived from different tests support the results reported here.

### 4.4. Conclusions

The current study demonstrates that resting-state functional connectivity with the hippocampus and the caudate nucleus can predict WM performance in people with MS. RSFC between the left caudate and right insula and between the left hippocampus and left SMG predicted better WM performance. RSFC between the left hippocampus and left vACC predicted worse WM performance. The hippocampal connections were also correlated with each other. Thus, we identify a caudate circuit and a potential hippocampal network whose aberrant activity at rest could potentially serve as a neural marker that signals impending WM impairment in people with MS. In a rehabilitation context, these brain markers could be combined with existing clinical tools to allow for earlier therapeutic efforts. By delivering targeted interventions *before* WM impairment is pronounced enough to be detected via neuropsychological assessment, we can further improve cognitive rehabilitation outcomes for people with MS.

## Acknowledgements

We would like to thank all research participants who made each of these studies possible.

## CRediT authorship contribution statement

**Christopher J. Cagna:** Conceptualization, Data curation, Formal analysis, Methodology, Validation, Visualization, Writing – original draft. **Ekaterina Dobryakova:** Conceptualization, Funding acquisition, Methodology, Resources, Supervision, Writing – review & editing. **Kai Sucich:** Formal analysis, Writing – review & editing. **Tien T. Tong:** Data curation, Software. **Joshua Sandry:** Conceptualization, Formal analysis, Funding acquisition, Methodology, Resources, Supervision, Writing – review & editing.

## References

Azzimonti, M., Preziosa, P., Pagani, E., Valasasina, P., Tedone, N., Vizzino, C., Rocca, M. A., & Filippi, M. (2023). Functional and structural MRI changes associated with cognitive worsening in multiple sclerosis: A 3-year longitudinal study (P11-3.003). Journal of Neurology, 270, 4296–4308. 10.1007/s00415-023-11778-z

Baddeley, A. D., & Hitch, G. (1974). Working memory. In The Psychology of Learning and Motivation (Bower, G. A., ed.) pp. 48–79. Academic Press.

Baddeley, A. (2000). The episodic buffer: A new component of working memory? Trends in Cognitive Sciences, 4(11), 417–423. 10.1016/S1364-6613(00)01538-2

Benedict, R. H. B., Schretlen, D., Groninger, L., & Brandt, J. (1998). The Hopkins verbal learning test-revised: Normative data and analysis of interform and test–retest reliability. Clinical Neuropsychologist, 12(4), 43–55. https://psycnet.apa.org/doi/10.1076/clin.12.1.43.1726

Borders, A. A., Ranganath, C., & Yonelinas, A. P. (2021). The hippocampus supports high-precision binding in visual working memory. Hippocampus, 32(3), 217–230. 10.1002/hipo.23401

Brissart, H., Morele, E., Baumann, C., & Debouverie, M. (2012). Verbal episodic memory in 426 multiple sclerosis patients: Impairment in encoding, retrieval, or both? Neurological Sciences, 33, 1117–1123. 10.1007/s10072-011-0915-7

Cagna, C. J., Ceceli, A. O., Sandry, J., Bhanji, J. P., Tricomi, E., & Dobryakova, E. (2022). Altered functional connectivity during performance feedback processing in multiple sclerosis. NeuroImage: Clinical, 37, 103287. 10.1016/j.nicl.2022.103287

Chiaravalloti, N. D., & DeLuca, J. (2008). Cognitive impairment in multiple sclerosis. The Lancet Neurology, 7(12), 1139–1151. 10.1016/S1474-4422(08)70259-X

Costers, L., Van Schependom, J., Laton, J., Baijot, J., Sjøgård, M., Wens, V., De Tiège, X., Goldman, S., D’Haeseleer, M., D’hooghe, M. B., Woolrich, M., & Nagels, G. (2020). The role of hippocampal theta oscillations in working memory impairment in multiple sclerosis. Human Brain Mapping, 42, 1376–1390. 10.1002/hbm.25299

Cowan, N. (1995). Attention and Memory: An Integrated Framework. In Attention and Memory: An Integrated Framework. Oxford University Press

DeLuca, J., Chiaravalloti, N. D., & Sandroff, B. M. (2020). Treatment and management of cognitive dysfunction in patients with multiple sclerosis. Nature Reviews Neurology, 16(6), 319–332. 10.1038/s41582-020-0355-1

Deschamps, I., Baum, S. R., & Gracco, V. L. (2014). On the role of the supramarginal gyrus in phonological processing and verbal working memory: Evidence from rTMS studies. Neuropsychologia, 53, 39–46. 10.1016/j.neuropsychologia.2013.10.015

Diedrichsen, J., Balsters, J. H., Flavell, J., Cussans, E., & Ramnani, N. (2009). A probabilistic MR atlas of the human cerebellum. NeuroImage, 46(1), 39–46. 10.1016/j.neuroimage.2009.01.045

Du, J., Rolls, E. T., Cheng, W., Li, Y., Gong, W., Qiu, J., & Feng, J. (2020). Functional connectivity of the orbitofrontal cortex, anterior cingulate cortex, and inferior frontal gyrus in humans. Cortex, 123, 185–199. 10.1016/j.cortex.2019.10.012

Duma, G. M., Mento, G., Cutini, S., Sessa, P., Baillet, S., Brigadoi, S., & Dell’Acqua, R. (2019). Functional dissociation of anterior cingulate cortex and intraparietal sulcus in visual working memory. Cortex, 121, 277–291. 10.1016/j.cortex.2019.09.009

Eijlers, A. J. C., Meijer, K. A., Wassenaar, T. M., Steenwijk, M. D., Uitdehaag, B. M. J., Barkhof, F., Wink, A. M., Geurts, J. J. G., & Schoonheim, M. M. (2017). Increased default-mode network centrality in cognitively impaired multiple sclerosis patients. Neurology, 88(10), 952–960. 10.1212/WNL.0000000000003689

Emch, M., von Bastian, C. C., & Koch, K. (2019). Neural correlates of verbal working memory: An fMRI meta-analysis. Frontiers in Human Neuroscience, 13:180. 10.3389/fnhum.2019.00180

Fox, M. D., & Greicius, M.(2010). Clinical applications of resting state functional connectivity. Frontiers in Systems Neuroscience, 4(19), 1–13. 10.3389/fnsys.2010.00019

Freedberg, M. V., Reeves, J. A., Fioriti, C. M., Murillo, J., Voss, J. L., & Wassermann, E. (2022). A direct test of competitive versus cooperative episodic-procedural network dynamics in human memory. Cerebral Cortex, 32, 4715–4732. 10.1093/cercor/bhab512

Freedberg, M. V. (2023). The balance of hippocampal and caudate network functional connectivity is associated with episodic memory performance and its decline across adulthood. Neuropsychologia, 191, 108723. 10.1016/j.neuropsychologia.2023.108723

Geurts, J. J. G., Bö, L., Roosendaal, S. D., Hazes, T., Daniëls, R., Barkhof, F., Witter, M. P., Huitinga, I., & van der Valk, P. (2007). Extensive hippocampal demyelination in multiple sclerosis. Journal of Neuropathology & Experimental Neurology, 66(9), 819–827. 10.1097/nen.0b013e3181461f54

Ghaziri, J., Tucholka, A., Girard, G., Boucher, O., Houde, J., Descoteaux, M., Obaid, S., Gilbert, G., Rouleau, I., & Nguyen, D. K. (2018). Subcortical structural connectivity of insular subregions. Scientific Reports, 8:8596. 10.1038/s41598-018-26995-0

Ghaziri, J., Fei, P., Tucholka, A., Obaid, S., Boucher, O., Rouleau, I., & Nguyen, D. K. (2024). Resting-state functional connectivity profile of insular subregions. Brain Sciences, 14(8), 742. 10.3390/brainsci14080742

González Torre, J. A., Cruz-Gómez, Á., J., Belenguer, A., Sanchis-Segura, C., Ávila, C., & Forn, C. (2017). Hippocampal dysfunction is associated with memory impairment in multiple sclerosis: A volumetric and functional connectivity study. Multiple Sclerosis Journal, 23(14), 1854–1863. 10.1177/1352458516688349

Grahn, J. A., Parkinson, J. A., & Owen, A. M. (2008). The cognitive functions of the caudate nucleus. Progress in Neurobiology, 86(3), 141–155. 10.1016/j.pneurobio.2008.09.004

Hulst, H. E., Schoonheim, M. M., Roosendaal, S. D., Popescu, V., Schweren, L. J. S., van der Werf, Y. D., Visser, L. H., Polman, C. H., Barkhof, F., & Geurts, J. J. G., (2012). Functional adaptive changes within the hippocampal memory system of patients with multiple sclerosis. Human Brain Mapping, 33, 2268–2280. 10.1002/hbm.21359

Hulst, H., Schoonheim, M. M., Van Geest, Q., Uitdehaag, B. M. J., Barkhof, F., & Geurts, J. J. G. (2015). Memory impairment in multiple sclerosis: Relevance of hippocampal activation and hippocampal connectivity. Multiple Sclerosis Journal, 21(13), 1705–1712. 10.1177/1352458514567727

Kaneda, M., & Osaka, N. (2008). Role of anterior cingulate cortex during semantic coding in verbal working memory. Neuroscience Letters, 436(1), 57–61. 10.1016/j.neulet.2008.02.069

Kumar, S., Joseph, S., Gander, P. E., Barascud, N., Halpern, A. R., & Griffiths, T. D. (2016). A brain system for auditory working memory. The Journal of Neuroscience, 36(16), 4492–4505. 10.1523/JNEUROSCI.4341-14.2016

Kurth, F., Zilles, K., Fox, P. T., Laird, A. R., & Eickhoff, S. B. (2010). A link between the systems: Functional differentiation and integration within the human insula revealed by meta-analysis. Brain Structure and Function, 214, 519–534. 10.1007/s00429-010-0255-z

Lenartowicz, A., & McIntosh, A. R. (2005). The role of anterior cingulate cortex in working memory is shaped by functional connectivity. Journal of Cognitive Neuroscience, 17(7), 1026–1042. 10.1162/0898929054475127

Leszczynski, M. (2011). How does the hippocampus contribute to working memory processing? Frontiers in Human Neuroscience, 5, 168. 10.3389/fnhum.2011.00168

Leszczynski, M., Fell, J., & Axmacher, N. (2015). Rhythmic working memory activation in the human hippocampus. Cell Reports, 13(6), 1272–1282. 10.1016/j.celrep.2015.09.081

Llorens, A., Bellier, L., Blenkmann, A. O., Endestad, T., Solbakk, A., & Knight, R. T. (2023). Decision and response monitoring during working memory are sequentially represented in the human insula. iScience, 26(10), 107653. 10.1016/j.isci.2023.107653

Menon, V., & Uddin, L. Q. (2010). Saliency, switching, attention and control: A network model of insula function. Brain Structure and Function, 214, 655–667. 10.1007/s00429-010-0262-0

Morfini, F., Whitfield-Gabrieli, S., & Nieto-Castañón, A. (2023). Functional connectivity MRI quality control procedures in CONN. Frontiers in Neuroscience, 17:1092125. 10.3389/fnins.2023.1092125

MS International Federation. (2025, April 10). What is MS?. https://www.msif.org/about-ms/what-is-ms/

Muhlert, N., Atzori, M., De Vita, E., Thomas, D. L., Samson, R. S., Wheeler-Kingshott, C. A. M., Geurts, J. J. G., Miller, D. H., Thompson, A. J., & Ciccarelli, O. (2014). Memory in multiple sclerosis is linked to glutamate concentration in grey matter regions. Journal of Neurology, Neurosurgery, & Psychiatry, 85, 833–839. 10.1136/jnnp-2013-306662

Müller, N. C. J., Konrad, B. N., Kohn, N., Muñoz-López, M., Czisch, M., Fernández, G., & Dresler, M. (2018). Hippocampal-caudate nucleus interactions support exceptional memory performance. Brain Structure and Function, 223, 1379–1389. 10.1007/s00429-017-1556-2

Nee, D. E., & Jonides, J. (2008). Neural correlates of access to short-term memory. Proceedings of the National Academy of Sciences, 105(37), 14228–14233. 10.1073/pnas.0802081105

Nee, D. E., & Jonides, J. (2011). Dissociable contributions of prefrontal cortex and the hippocampus to short-term memory: Evidence for a 3-state model of memory. NeuroImage, 54(2), 1540–1548. 10.1016/j.neuroimage.2010.09.002

Papadopoulos, D., Dukes, S., Patel, R., Nicholas, R., Vora, A., Reynolds, R. (2009). Substantial archaeocortical atrophy and neuronal loss in multiple sclerosis. Brain Pathology, 19(2), 238–253. 10.1111/j.1750-3639.2008.00177.x

Parkes, L., Fulcher, B., Yücel, M., & Fornito, A. (2018). An evaluation of the efficacy, reliability, and sensitivity of motion correction strategies for resting-state functional MRI. NeuroImage, 171, 415–436. 10.1016/j.neuroimage.2017.12.073

Pitteri, M., Galazzo, I. B., Brusini, L., Cruciani, F., Dapor, C., Marastoni, D., Menegaz, G., & Calabrese, M. (2021). Microstructural MRI correlates of cognitive impairment in multiple sclerosis: The role of deep gray matter. Diagnostics, 11(6), 1103. 10.3390/diagnostics11061103

Purcell, J., Rapp, B., Martin, R. C. (2021). Distinct neural substrates support phonological and orthographic working memory: Implications for theories of working memory. Frontiers in Neurology, 12:681141. 10.3389/fneur.2021.681141

Qi, L., Rosenberg, M. D., Yoo, K., Hsu, T. W., O’Connell, T. P., Chun, M. M. (2018). Resting-state functional connectivity predicts cognitive impairment related to Alzheimer’s Disease. Frontiers in Aging Neuroscience, 10(94), 1–10. 10.3389/fnagi.2018.00094

Rocca, M. A., Valsasina, P., Absinta, M., Riccitelli, G., Rodegher, M. E., Misci, P., Rossi, P., Falini, A., & Comi, G. (2010). Default-mode network dysfunction and cognitive impairment in progressive MS. Neurology, 74(16), 1252–1259. 10.1212/WNL.0b013e3181d9ed91

Rocca, M. A., Schoonheim, M. M., Valsasina, P., Geurts, J. J. G., & Filippi, M. (2022). Task- and resting-state fMRI studies in multiple sclerosis: From regions to systems and time-varying analysis. Current status and future perspective. NeuroImage: Clinical, 35, 103076. 10.1016/j.nicl.2022.103076

Roosendaal, S. D., Moraal, B., Vrenken, H., Castelijns, J. A., Pouwels, P. J. W., Barkhof, F., & Geurts, J. J. G. (2008). In vivo MR imaging of hippocampal lesions in multiple sclerosis. Journal of Magnetic Resonance Imaging, 27(4), 726–731. 10.1002/jmri.21294

Sacco, R., Bisecco, A., Corbo, D., Della Corte, M., d’Ambrosio, A., Docimo, R., Gallo, A., Esposito, F., Esposito, S., Cirillo, M., Lavorgna, L., Tedeschi, G., & Bonavita, S. (2015). Cognitive impairment and memory disorders in relapsing-remitting multiple sclerosis: The role of white matter, gray matter, and hippocampus. Journal of Neurology, 262, 1691–1697. 10.1007/s00415-015-7763-y

Sandry, J. & Sumowski, J. F. (2014). Working memory mediates the relationship between intellectual enrichment and long-term memory in multiple sclerosis: An exploratory analysis of cognitive reserve. Journal of the International Neuropsychological Society, 20, 868 – 872. 10.1017/S1355617714000630

Sandry, J. (2015). Working memory and memory loss in neurodegenerative disease. Neurodegenerative Disease Management. 5(1), 1–4. 10.2217/nmt.14.51

Sumowski, J. F., Rocca, M. A., Leavitt, V. M., Riccitelli, G., Sandry, J., DeLuca, J., Comi, G., & Filippi, M. (2016). Searching for the neural basis of reserve against memory decline: Intellectual enrichment linked to larger hippocampal volume in multiple sclerosis. European Journal of Neurology, 23, 39–44. 10.1111/ene.12662

Schoonheim, M. M., Geurts, J. J. G., & Barkhof, F. (2010). The limits of functional reorganization in multiple sclerosis. Neurology, 74(16), 1246–1247. 10.1212/WNL.0b013e3181db9957

Schoonheim, M. M., Meijer, K. A., & Geurts, J. J. G. (2015). Network collapse and cognitive impairment in multiple sclerosis. Frontiers in Neurology, 6:82. 10.3389/fneur.2015.00082

Smith, R., Lane, R. D., Alkozei, A., Bao, J., Smith, C., Sanova, A., Nettles, M., & Killgore, W. D. S. (2018). The role of the medial prefrontal cortex in the working memory maintenance of one’s own emotional responses. Scientific Reports, 8, 3460. 10.1038/s41598-018-21896-8

Sumowski, J. F., Leavitt, V. M., Rocca, M. A., Inglese, M., Riccitelli, G., Buyukturkoglu, K., Meani, A., & Filippi, M. (2017). Mesial temporal lobe and subcortical grey matter volumes differentially predict memory across stages of multiple sclerosis. Multiple Sclerosis Journal, 24(5), 675–678. 10.1177/1352458517708873

Tsouki, F., & Williams, A. (2021). Multifaceted involvement of microglia in gray matter pathology in multiple sclerosis. Stem Cells, 2021, 1–15. 10.1002/stem.3374

Tulving, E., (2002). Episodic memory: From mind to brain. Annual Review of Psychology, 53, 1–25. 10.1146/annurev.psych.53.100901.135114

Unsworth, N., & Engle, R. W. (2006). Simple and complex memory spans and their relation to fluid abilities: Evidence from list-length effects. Journal of Memory and Language, 54(1), 68–80. 10.1016/j.jml.2005.06.003

Venkataraman, A., Whitford, T. J., Westin, C., Golland, P., & Kubicki, M. (2012). Whole brain resting state functional connectivity abnormalities in schizophrenia. Schizophrenia Research, 139(1-3), 7–12. 10.1016/j.schres.2012.04.021

Venkatesan, U. M., Dennis, N. A., & Hillary, F. G. (2015). Chronology and chronicity of altered resting-state functional connectivity after traumatic brain injury. Journal of Neurotrauma, 32(4), 252–264. 10.1089/neu.2013.3318

Wechsler, D. (2008). WAIS-IV Administration and Scoring Manual. Psychological Corporation, San Antonio, TX.

Whitfield-Gabrieli, S., & Nieto-Castañón, A. (2012). *CONN:* A functional connectivity toolbox for correlated and anticorrelated brain networks. Brain connectivity, 2(3), 125–141. 10.1089/brain.2012.0073

Wisch, J. K., Roe, C. M., Babulal, G. M., Schindler, S. E., Fagan, A. M., Benzinger, T. L., Morris, J. C., & Ances, B. M. (2020). Resting state functional connectivity signature differentiates cognitively normal from individuals who convert to symptomatic Alzheimer Disease. Journal of Alzheimer’s Disease, 74(4), 1085–1095. 10.3233/JAD-191039

Yonelinas, A., Hawkins, C., Abovian, A., & Aly, M. (2024). The role of recollection, familiarity, and the hippocampus in episodic and working memory. Neuropsychologia, 193, 108777. 10.1016/j.neuropsychologia.2023.108777

Zhang, J., Kucyi, A., Raya, J., Nielsen, A. N., Nomi, J. S., Damoiseaux, J. S., Greene, D. J., Horovitz, S. G., Uddin, L. Q., & Gabrieli-Whitfield, S. (2021). What have we really learned from functional connectivity in clinical populations? NeuroImage, 242, 118466. 10.1016/j.neuroimage.2021.118466

Zuppichini, M. D., & Sandry, J. (2018). Pilot investigation of the relationship between hippocampal volume and pattern separation deficits in multiple sclerosis. Multiple Sclerosis and Related Disorders, 26, 157–163. 10.1016/j.msard.2018.09.016

