## Supplementary Material for "Resting-state Functional Connections with the Hippocampus and with the Caudate Nucleus Predict Working Memory Performance in Multiple Sclerosis"

**Table S1.**
Correlations (Pearson's $r$) for significant right hippocampal resting-state functional connections and working memory composite scores in the MS group.

#### MS Group

#### Seed: Right Hippocampus

| | Variable | 1 | 2 | 3 | 4 | 5 | 6 | 7 | 8 | 9 | 10 | 11 |
|---|---|---|---|---|---|---|---|---|---|---|---|---|
| 1 | WM Composite | --- | | | | | | | | | | |
| 2 | R Hipp | .07 | --- | | | | | | | | | |
| 3 | L Hipp | .24 | .71*** | --- | | | | | | | | |
| 4 | SFG | -.06 | -.56*** | -.43** | --- | | | | | | | |
| 5 | L SMG | .09 | -.37* | -.18 | .37* | --- | | | | | | |
| 6 | Supracalcarine Cortex | .06 | -.67*** | -.46** | .51*** | .67*** | --- | | | | | |
| 7 | R Primary Visual Cortex | -.16 | -.57*** | -.53*** | .46** | .38* | .64*** | --- | | | | |
| 8 | L dlPFC | .02 | -.29 | -.13 | .36* | .43** | .42** | .44** | --- | | | |
| 9 | L Premotor Area | -.20 | -.54*** | -.57*** | .58*** | .40** | .38* | .45** | .29 | --- | | |
| 10 | R SMG | -.27 | -.31* | -.21 | .27 | .27 | .36* | .26 | .17 | .24 | --- | |
| 11 | L Pars Orbitalis | -.15 | .03 | -.06 | .20 | .17 | .01 | .22 | .37* | -.002 | .14 | --- |

*Notes*: N = 26. WM = working memory (i.e., working memory composite measure); R = Right; L = Left; Hipp = Hippocampus; SFG = Superior Frontal Gyrus; SMG = Supramarginal Gyrus; dlPFC = Dorsolateral Prefrontal Cortex. *$p$ < .05; **$p$ < .01; ***$p$ < .001 (uncorrected).

**Table S2.**
Correlations (Pearson's *r*) for significant right hippocampal resting-state functional connections and working memory composite scores in the HC group.

#### HC Group

##### Seed: Right Hippocampus

| | Variable | 1 | 2 | 3 | 4 | 5 | 6 |
|---|---|---|---|---|---|---|---|
| 1 | WM Composite | --- | | | | | |
| 2 | R Hipp | .04 | --- | | | | |
| 3 | L Hipp | .23 | .69*** | --- | | | |
| 4 | R vPCC | -.24 | .21 | .11 | --- | | |
| 5 | R PMA | .43 | -.36* | -.37* | -.25 | --- | |
| 6 | L Parahipp | -.20 | .62*** | .42** | .18 | -.32 | --- |

*Notes*: N = 15. WM = working memory (i.e., working memory composite measure); R = Right; L = Left; Hipp = Hippocampus; vPCC = Ventral Posterior Cingulate Cortex; PMA = Premotor Area; Parahipp = Parahippocampus. *$p < .05$; **$p < .01$; ***$p < .001$ (uncorrected).

**Table S3.**
Correlations (Pearson's $r$) for significant left hippocampal resting-state functional connections and working memory composite scores in the MS group.

#### MS Group

##### Seed: Left Hippocampus

| | Variable | 1 | 2 | 3 | 4 | 5 | 6 | 7 | 8 | 9 | 10 | 11 |
|---|---|---|---|---|---|---|---|---|---|---|---|---|
| 1 | WM Composite | --- | | | | | | | | | | |
| 2 | L Hipp | .02 | --- | | | | | | | | | |
| 3 | R Hipp | .09 | .64*** | --- | | | | | | | | |
| 4 | R Precuneus | .10 | -.56*** | -.50*** | --- | | | | | | | |
| 5 | R SMG | .003 | -.28 | -.17 | .37* | --- | | | | | | |
| 6 | L aPFC | .08 | .36* | .23 | -.29 | -.50*** | --- | | | | | |
| 7 | L vACC | -.49** | .44** | .33** | -.47** | -.43** | .38* | --- | | | | |
| 8 | L dlPFC | -.04 | -.17 | -.22 | .38* | .39* | -.10 | -.10 | --- | | | |
| 9 | R dlPFC | .12 | -.21 | -.19 | .31* | .33* | -.10 | -.18 | .51*** | --- | | |
| 10 | R LOC | .17 | -.45** | -.32* | .61*** | .37* | -.09 | -.33* | .35* | .30 | --- | |
| 11 | L SMG | .49* | -.22 | -.15 | .47** | .44** | -.12 | -.45** | .30 | .33* | .30 | --- |

*Notes*: N = 26. WM = working memory (i.e., working memory composite measure); R = Right; L = Left; Hipp = Hippocampus; SMG = Supramarginal Gyrus; aPFC = Anterior Prefrontal Cortex; vACC = Ventral Anterior Cingulate Cortex; dlPFC = Dorsolateral Prefrontal Cortex; LOC = Lateral Occipital Cortex. *$p < .05$; **$p < .01$; ***$p < .001$ (uncorrected). Benjamini-Hochberg procedures (False Discovery Rate $\alpha = .05$) were employed to correct for the number of tests conducted with the WM composite measure (see Methods). Corrected p values for significant associations with WM appear in the Results.

**Table S4.**
Correlations (Pearson's *r*) for significant left hippocampal resting-state functional connections and working memory composite scores in the HC group.

#### HC Group

##### Seed: Left Hippocampus

| | Variable | 1 | 2 | 3 | 4 |
|---|---|---|---|---|---|
| 1 | WM Composite | --- | | | |
| 2 | L Hipp | -.16 | --- | | |
| 3 | R Hipp | -.11 | .70*** | --- | |
| 4 | L Temporal Pole | -.08 | .64*** | .50** | --- |

*Notes*: N = 15. WM = working memory (i.e., working memory composite measure); R = Right; L = Left; Hipp = Hippocampus; *$p < .05$; **$p < .01$; ***$p < .001$ (uncorrected).

**Table S5.**
Correlations (Pearson's *r*) for significant right caudal resting-state functional connections and working memory composite scores in the MS group.

### MS Group

#### Seed: Right Caudate

| | Variable | 1 | 2 | 3 | 4 |
|---|---|---|---|---|---|
| 1 | WM Composite | --- | | | |
| 2 | R+L Caudate | -.07 | --- | | |
| 3 | R Parahipp | -.15 | .51*** | --- | |
| 4 | L dlPFC | -.03 | .45** | .60*** | --- |

*Notes*: N = 26. WM = working memory (i.e., working memory composite measure); R = Right; L = Left; Parahipp = Parahippocampus; dlPFC = Dorsolateral Prefrontal Cortex. **p* < .05; ***p* < .01; ****p* < .001 (uncorrected).

**Table S6.**
Correlations (Pearson's *r*) for significant right caudal resting-state functional connections and working memory composite scores in the HC group.

### HC Group

#### Seed: Right Caudate

| | Variable | 1 | 2 | 3 |
|---|---|---|---|---|
| 1 | WM Composite | --- | | |
| 2 | R+L Caudate | -.30 | --- | |
| 3 | R Midbrain | .11 | .25 | --- |

*Notes*: N = 15. WM = working memory (i.e., working memory composite measure); R = Right; L = Left.

**Table S7.**
Correlations (Pearson's *r*) for significant left caudal resting-state functional connections and working memory composite scores in the MS group.

#### MS Group

#### Seed: Left Caudate

| | Variable | 1 | 2 | 3 | 4 | 5 | 6 | 7 | 8 | 9 |
|---|---|---|---|---|---|---|---|---|---|---|
| 1 | WM Composite | --- | | | | | | | | |
| 2 | R+L Caudate | -.05 | --- | | | | | | | |
| 3 | R vPCC | -.08 | -.30 | --- | | | | | | |
| 4 | R Cerebellar Lobule VIIb | .05 | -.59*** | .49*** | --- | | | | | |
| 5 | L Cerebellar Crus I | .09 | -.58*** | .48*** | .66*** | --- | | | | |
| 6 | R Insula | .48* | .41** | -.38* | -.53*** | -.36* | --- | | | |
| 7 | R ITG | -.02 | -.27 | .48*** | .52*** | .54*** | -.39** | --- | | |
| 8 | R Cerebellar Crus II | .02 | -.51*** | .38* | .58*** | .59*** | -.33* | .41** | --- | |
| 9 | L Cerebellar Region VI | -.11 | -.36* | .37* | .45** | .46** | -.30 | .40** | .38* | --- |

*Notes*: N = 26. WM = working memory (i.e., working memory composite measure); R = Right; L = Left; vPCC = Ventral Posterior Cingulate Cortex; ITG = Inferior Temporal Gyrus. *$p$ < .05; *$p$ < .01; ***$p$ < .001 (uncorrected). Benjamini-Hochberg procedures (False Discovery Rate α = .05) were employed to correct for the number of tests conducted with the WM composite measure (see Methods). Corrected p values for significant associations with WM appear in the Results.

**Table S8.**
Correlations (Pearson's *r*) for significant left caudal resting-state functional connections and working memory composite scores in the HC group.

### HC Group

#### Seed: Left Caudate

| | Variable | 1 | 2 |
|---|---|---|---|
| 1 | WM Composite | --- | |
| 2 | R+L Caudate | -.25 | --- |

*Notes*: N = 15. WM = working memory (i.e., working memory composite measure); R = Right; L = Left.
